# Developmental Dynamics of Human Cardiogenesis: A multi-omic reference and its disruption in Trisomy 21

**DOI:** 10.1101/2024.04.29.591736

**Authors:** James Cranley, Kazumasa Kanemaru, Semih Bayraktar, Vincent Knight-Schrijver, Rebecca Hulbert, Eva Lana-Elola, Rifdat Aoidi, Jan Patrick Pett, Anna Wilbrey-Clark, Krzysztof Polanski, Monika Dabrowska, Ilaria Mulas, Harriet Johnson, Laura Richardson, Claudia I. Semprich, Rakeshlal Kapuge, Shani Perera, Xiaoling He, Siew Yen Ho, Nadav Yayon, Liz Tuck, Kenny Roberts, Jack A. Palmer, Hongorzul Davaapil, Laure Gambardella, Minal Patel, Richard Tyser, Andreia Sofia Bernardo, Victor L. J. Tybulewicz, Sanjay Sinha, Sarah A. Teichmann

## Abstract

Developmental dynamics involve the specification of diverse cell types and their spatial organization into multicellular niches. Here, we combine single-cell and spatial multiomics to define 19 distinct tissue niches in the developing heart, leading to the development of a context-aware, resolution-agnostic niche classification tool (`TissueTypist`). Applying high-resolution spatial profiling to the developing sinoatrial node, we resolve three pacemaker cell subtypes arrayed along a linear axis. First trimester subpopulations, such as the pacemaker cells in the sinus horn and sinoatrial node head region, display neuro-attractant programmes and interact with parasympathetic neurons via interactions including Semaphorin-Plexin signalling. Temporal trajectories map maturation of atrial and ventricular cardiomyocytes, uncovering a lipid-metabolic switch and potential key regulators of cell type identity. In the ventricle, we identify cellular and transcriptional gradients along both pseudotime and transmural axes, offering new molecular insights into myocardial compaction and maturation. Comparative profiling of euploid and trisomy 21 hearts shows a depletion of compact cardiomyocytes and heightened apoptosis, validated in isogenic-matched trisomy 21 and euploid iPSC-derived cardiomyocytes. This implicates disrupted myocardial growth may be a mechanism for Down’s syndrome-associated congenital heart disease. Overall, we deliver a spatially resolved framework of human cardiac development, enabling systematic exploration of developmental niches in health and disease.

## 1 Introduction

Human cardiogenesis requires coordinated interactions among diverse cell types originating from cardiogenic mesoderm and neural crest. These populations orchestrate key processes such as the formation of the sinoatrial node (SAN), the primary pacemaker of the heart, and cardiomyocyte maturation [1**?**]. This intricate choreography is highly sensitive to genetic, epigenetic, and environmental perturbations, with disruptions linked to congenital heart disease (CHD)—a heterogeneous set of malformations affecting 1% of live births and contributing to approximately 4% of infant mortality [2]. The ∼50-fold increased incidence of CHD in trisomy 21 highlights the vulnerability of cardiac development to altered gene dosage [3]. Many known risk genes for CHD are transcription factors [4], and their mutations often produce pleiotropic phenotypes, suggesting disruption of complex, spatially and temporally governed gene regulatory networks (GRNs) that orchestrate morphogenesis across the developing heart. Despite significant advancements in the identification of causal genes, more than half of CHD cases remain without a definitive genetic diagnosis [5]. Additionally, the cellular mechanisms contributing to CHD are not yet fully characterised.

While human induced pluripotent stem cell-derived cardiomyocytes (hiPSC-CMs) have provided insight into aspects of cardiomyocyte identity and maturation, they imperfectly model *in vivo* development and often fail to recapitulate key features of tissue organization or cellular diversity [6]. Recent efforts to profile the foetal heart using transcriptomic and spatial ‘-omics’ approaches have revealed spatial gene expression patterns and cell-type heterogeneity [7–9]. However, these studies have generally been limited in developmental coverage, spatial resolution, or modality of data types. Furthermore, robust methodologies for identifying and annotating the cellular niches within the developing heart, utilising varying resolutions of spatial omics data, have yet to be established. Consequently, there remains a clear need for comprehensive, multimodal, and spatially resolved atlases that elucidate the cellular and molecular dynamics of human cardiac development.

To address these gaps, we constructed a spatially-resolved multi-modal reference atlas of the developing human, spanning 4 to 20 post-conceptional weeks (PCW). This spatial atlas integrates single-cell transcriptomic references from our accompanying study [10], while incorporating paired single-nucleus ATAC-seq data, facilitating comprehensive multiomic analysis. This atlas delineates 19 transcriptionally distinct tissue domains using 4̃0,000 spatial transcriptomic pixels. To leverage the identified tissue domains, we introduce `TissueTypist`, a new computational tool for cellular niche identification in spatial transcriptomic data of differing resolutions. We show that its context-aware design gives improved accuracy, and we benchmark it on both in-house and public datasets. Using the `OrganAxis` framework [11] we overcome differences in sample size by constructing common coordinate axes, including a transmural axis to map cell types and compaction in the ventricular wall, and a trans-SAN axis to localise pacemaker cell subtypes and their associated parasympathetic inputs. Finally, by integrating age-matched euploid and trisomy 21 datasets, we identify a depletion of a specific ventricular cardiomyocyte subtype in trisomy 21 hearts. We link this to increased apoptosis in proliferating cardiomyocytes. We validate this observation using an isogenic-matched trisomy 21 hiPSC-CM model. Together this sheds light on the mechanisms underlying CHD in Down syndrome.

Overall, this study offers a foundational resource and analytical framework for decoding human cardiac development, including its cellular niches and the perturbations associated with congenital disorders.

## 2 Results

We collected 24 human euploid foetal hearts spanning 4 to 20 post-conception weeks (PCWs), which were subjected to either single-cell RNA sequencing, single-nucleus Multiome sequencing (paired RNA- and ATAC-seq), or spatial transcriptomics (**Fig. 1A**, **Supp. Fig. S1A**, **Supp. Table 1-2**). Following quality control of single-cell and single-nucleus RNA-seq data, we retained 297,473 cells and nuclei with high-quality gene expression profiles (**Supp. Fig. S1A**). We incorporated spatial transcriptomics data from 8 hearts, totalling 32 capture areas, to further enrich the dataset. Spatial transcriptome profiling was primarily conducted using 10x Genomics Visium, with a subset of samples profiled at higher resolution using Visium HD and Xenium, enabling detailed cellular and molecular localisation.

**Fig. 1:**
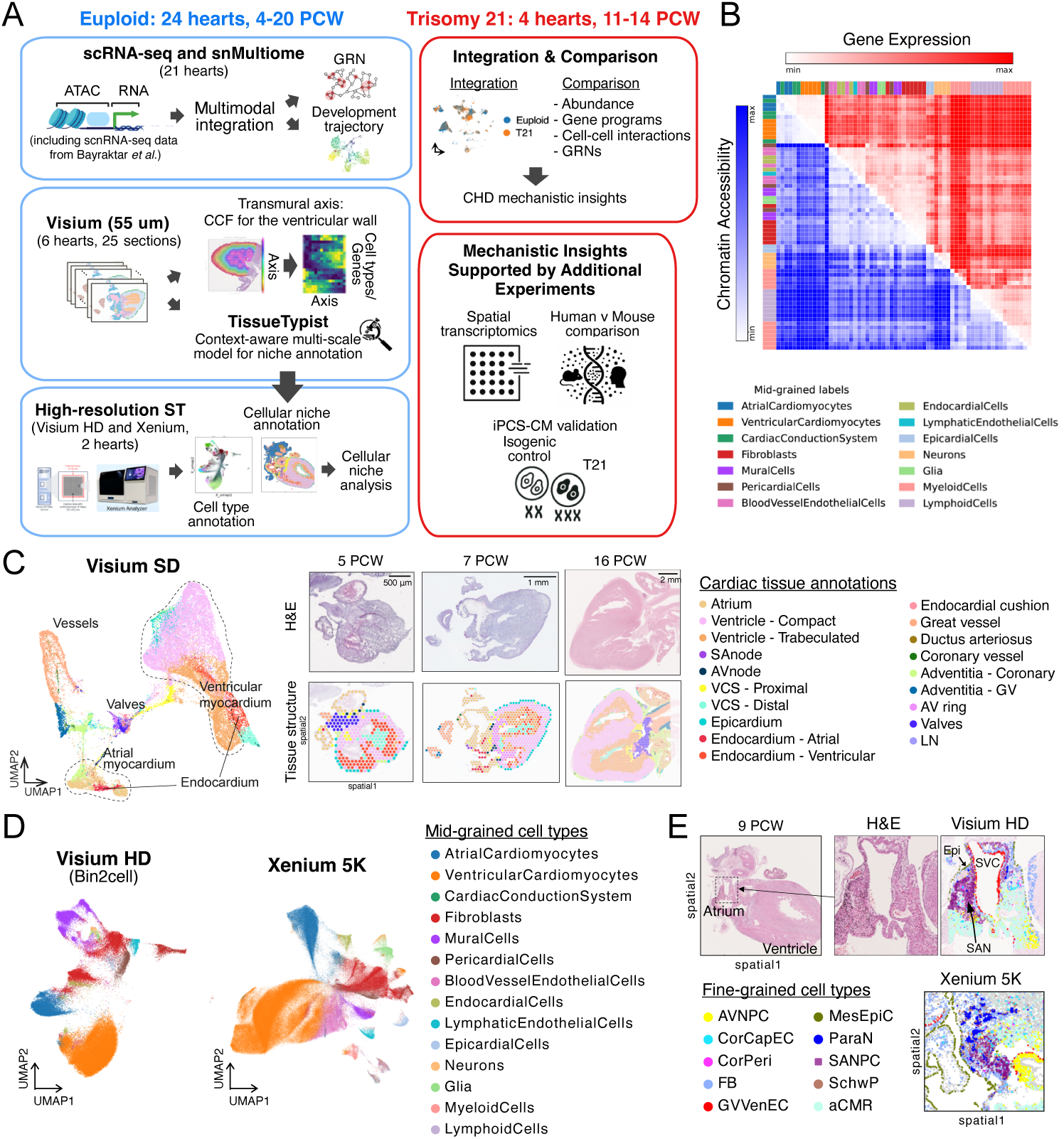
Multiomic atlas of the developing human heart. **A:** Overview of study design, data modalities, and analysis strategies. sc/snRNA-seq [**?**], Multiome, and spatial transcriptomics data were generated from a total of 24 human euploid fetal hearts between the ages of 4 and 20 PCW. Euploid datasets and models were integrated with the newly generated Trisomy 21 datasets (between the ages of 11 and 14 PCW) for comparative analysis. Additional experiments were performed to evaluate mechanistic insights. **B:** Heatmap showing Euclidean distance matrices of fine-grained cell types in gene expression (upper triangle, red) and chromatin accessibility (lower triangle, blue) spaces. Distances are summarised as the median distance for the cells in each fine-grained cell type. Each row and column is a fine-grained cell type. Row and column edge colours represent the parent mid-grained cell type category. The Pearson correlation co-efficient between these 2 distance matrixes was 0.76 with a p-value of 0.01 (Mantel test). **C:** UMAP embedding of estimated cell type abundances (`cell2location`) of the Visium SD spots across 22 spatial sections (39,563 spots). Spots were annotated based on clustering, estimated abundance of cell types in each cluster, and histological structure as defined by expert annotations using hematoxylin and eosin (H&E) images (Methods). Representative whole heart Visium sections with structural annotations, are shown on the right. **D:** UMAP embedding of gene expression data of the single-cell level high-resolution spatial transcriptomics: Visium HD (129,803 cells) and Xenium-5K (362,277 cells). Cell segmentation for each modality was performed using `bin2cell` or Xenium *in situ* multimodal cell segmentation algorithm, respectively. **E:** Representative fine-grained cell type localisations in the SAN of the Visium HD and Xenium-5K data. **Abbreviations**: PCW – post-conception weeks; sc/snRNA-seq – single-cell/single-nucleus RNA sequencing; ATAC – assay for transposase-accessible chromatin; SD – standard definition (55 µm spot diameter); UMAP – Uniform Manifold Approximation and Projection; H&E – hematoxylin and eosin. The abbreviations of the fine-grained cell types are listed in Supplementary Table 6.

Cell-type annotations were transferred from our companion study [10], which defines a three-level hierarchical system: 6 coarse-grained labels, 14 mid-grained labels, and 63 fine-grained labels (**Supp. Fig. S1A**). Of the nuclei subjected to Multiome sequencing, 167,022 from 11 hearts (spanning 4 to 20 PCW) also passed ATAC-seq quality control (**Supp. Fig. S1B**) with cell-type annotations inherited from their RNA-derived labels. Chromatin-accessible regions (peaks) were identified for each finegrained cell type, and their union resulted in a total of 508,040 peaks. 66% of these peaks overlapped—defined as more than 100 base pairs—with the ENCODE candidate cis-regulatory elements (Registry V3) [12]. We observed strong concordance between ATAC and RNA profiles: the transcriptionally defined cell types were distinctly separated in the ATAC UMAP embedding (**Supp. Fig. S1B**), and a Mantel test comparing inter-cell-type distance matrices between RNA and ATAC modalities revealed a strong correlation (r = 0.79; **Fig. 1B**), supporting the concordance between transcriptional and chromatin accessibility landscapes.

### 2.1 Spatial transcriptomics reveal multicellular niches

To explore cellular niches in the developing human heart, we first mapped our cell type annotations to spatial transcriptomic data using `cell2location` [13], inferring the abundance of each cell type per spot (**Supp. Fig. S1A**). We then integrated the spot-by-cell type abundance data (39,563 spots) from 22 Visium SD (Standard Definition, 55 µm resolution) sections (**Fig. 1C**). These spots were annotated with tissue structures based on clustering, estimated abundance of cell types in each cluster, and histological structure as defined by expert annotations using the associated H&E images (**Fig. 1C**). This analysis revealed shared structural gene expression signatures across chronological age, which supports the overall consistency of our spatial dataset (**Fig. 1C**).

To explore the localisation of cells and genes, we conducted high-resolution spatial transcriptomics utilising both sequencing-based (Visium HD, 2 µm resolution) and imaging-based (Xenium) technologies (9 PCW and 16 PCW hearts). The 2 µm bin data of the Visium HD output was aggregated into cells using the `bin2cell` algorithm [14]. This yielded rich profiles (median number of genes: 1138, median total counts: 2601) for 129,803 Visium HD cells (**Fig. 1D**). The Multimodal morphology-based cell segmentation algorithm (10x Genomics) was applied to Xenium datasets (5K Human Pan Tissue and Pathways Panel), which yielded 362,277 cells (**Fig. 1D**). We annotated cell types in both technologies through a combination of label transfer from reference single-cell/nucleus data (`CellTypist`[15]) and marker-driven manual annotation. The identified cell types demonstrated a high degree of consistency in comparison to those identified in the single-cell and nucleus data (**Supp. Fig. S1C**). Both sections utilised for the Visium HD and Xenium 5K assays included sinoatrial node (SAN) tissues. SAN pacemaker cells were found to be in close proximity to various cell types, including neural cells, coronary cells, fibroblasts, and cells derived from the developing superior vena cava (great vessel venous endothelial cells: GVVenEC) (**Fig. 1E**).

Each annotated tissue structure in the 55 µm resolution Visium dataset showed a distinct combination of cell-type abundances, reflecting unique multicellular niches (**Fig. 2A**). The SAN showed enrichment of pacemaker cells, as well as CX3CR1+ macrophages and parasympathetic neurons. This is consistent with the observation in the post-natal human heart that nodal autonomic innervation is predominantly vagal [16, 17] (**Fig. 2A**). LYVE1+ tissue-resident macrophages were highly abundant in vessel-associated structures (adventitia of coronary and great vessels) and also in the epicardium. The ventricular compact myocardium showed a high abundance of coronary vascular cells, such as capillary endothelial cells and pericytes (**Fig. 2A**). The adventitia of the great vessels, unlike coronary vessels, was highly enriched for neuron progenitors. This finding is consistent with our companion study, which demonstrates higher neuroattractant gene signatures in smooth muscle cells of great vessels compared to those of coronary vessels [10]. Since parasympathetic innervation is known to occur first, these may represent sympathetic precursor cells that are part of the developing plexi growing along the arterial surfaces [18] (**Fig. 2A**). We found a higher abundance of parasympathetic compared to sympathetic neurons across various tissues, including the node, atrium, and ventricle, particularly at earlier stages (**Supp. Fig. S2A**). In single-nucleus RNA sequencing data [10], parasympathetic and sympathetic neurons were predominantly detected in samples dissected from the atria, and sympathetic neurons were also enriched in samples dissected from the aorta (**Supp. Fig. S2B**). These findings suggest that ganglionated plexi are primarily localised in the atrial tissues and, notably, parasympathetic axons begin to innervate broader regions of the heart early in development (4-5 PCW). This is in keeping with previous animal studies which suggest parasympathetic innervation begins in the first trimester and precedes sympathetic innervation [18].

**Fig. 2:**
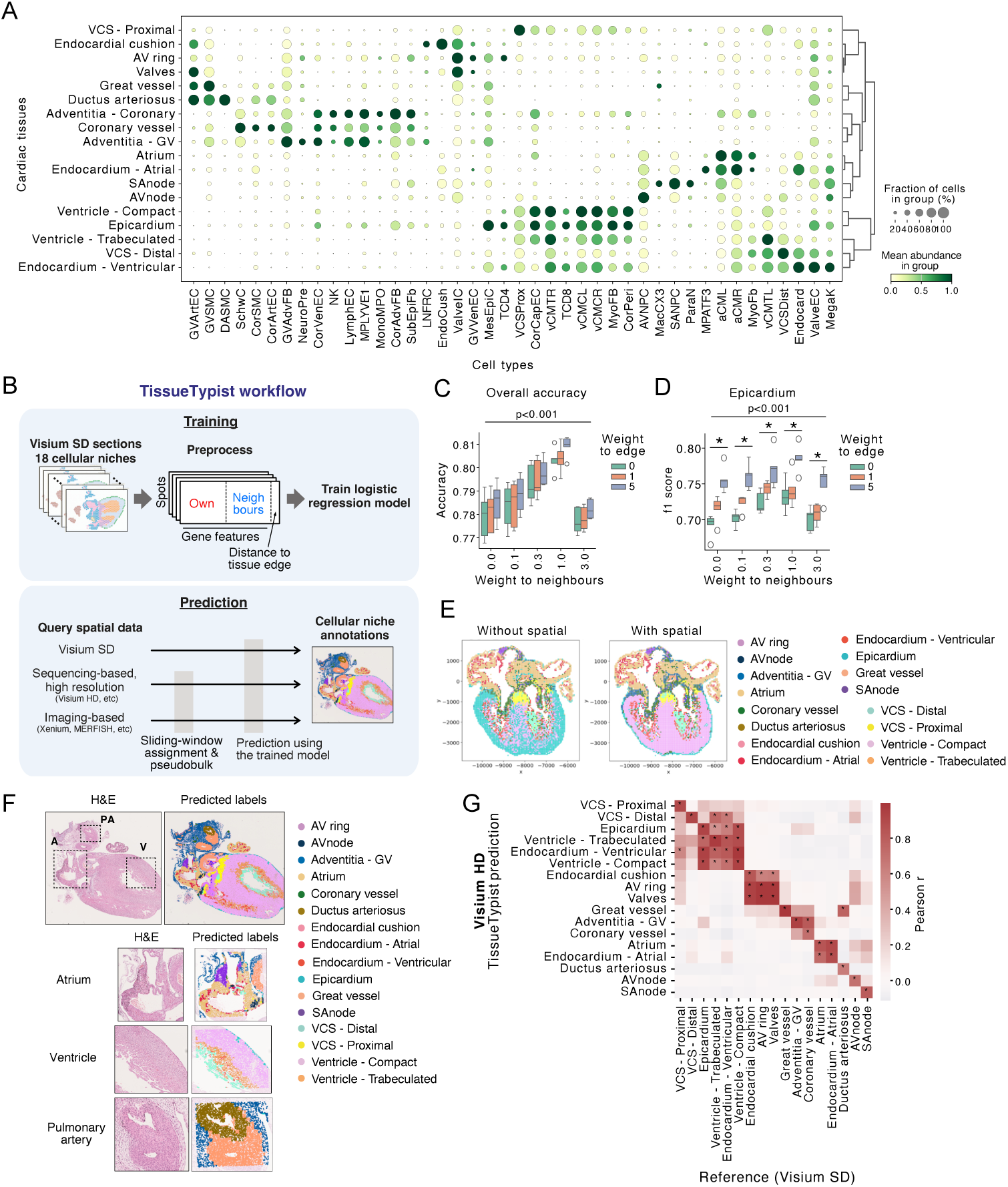
Spatial transcriptomics reveals multicellular niches. **A:** Dotplot displaying the estimated fine-grained cell (columns) abundance per tissue structure annotation (rows). Summarised abundances were normalised for each cell type. **B:** Workflow of the TissueTypist model training and prediction. The training workflow employs a context-aware model, integrating each spot’s gene expression profile with information from neighbouring spots and its distance to the tissue edge (upper box). The prediction workflow delivers rapid results across various spatial transcriptomics platforms operating at different resolution scales, utilising a sliding-window and pseudobulk assignment approach (lower box). **C, D:** Cross-validated overall accuracy (C) and F1 (D) scores for the epicardium niche across different weightings of neighbouring-spot information and distance-to-edge features. F1 score is the harmonic mean of precision and recall, providing a balanced measure that considers both false positives and false negatives. **E, F:** Prediction results on an external imaging–based spatial transcriptomics (MERFISH) dataset [9], comparing models with and without spatial context (E), and in-house Visium HD data of 9 PCW heart (F). **G:** Correlations of cell-type proportions per niche between the query Visium HD data and reference datasets. **Abbreviations**: Visium SD – Visium Standard Definition (55 *µ*m); H&E – hematoxylin and eosin. The abbreviations of the fine-grained cell types are listed in Supplementary Table 6.

### 2.2 TissueTypist - a context-aware multi-resolution model for niche annotation in spatial transcriptomic data

A total of 19 cardiac tissues were annotated, including seven myocardial and five vascular structures. Having defined this set of tissue structures, we developed a logistic regression classifier, “`TissueTypist`” (https://github.com/Teichlab/TissueTypist), trained not only on the intrinsic transcriptomic signatures of each tissue but also on spatial contextual features: the transcriptomic profiles of neighbouring regions and the distance to the tissue edge (context-aware model), (**Fig. 2B, Supp. Fig. 2C**)). Furthermore, we designed the prediction workflow to be compatible with datasets of higher spatial resolution, such as Visium HD, Xenium, and MERFISH, by incorporating a multi-scale modelling approach (multi-resolution model, **Fig. 2B**). We demonstrate that incorporating local features significantly improves tissue type prediction accuracy across most tissue types, with particularly pronounced benefits in specific tissues (**Fig. 2C, Supp. Fig. S2D**). The distance to the tissue edge proved especially advantageous for the prediction accuracy of the “epicardium” niche (**Fig. 2D**). Evaluation using a public high-resolution spatial transcriptomics dataset [9] also indicated that the context-aware model enhances the accuracy of prediction outputs (**Fig. 2E**). By applying the model to Visium HD 8 *µ*m-bin data, we recovered tissue annotations, including SAN, compact and trabeculated myocardium, and ductus arteriosus, without the need for cell segmentation and cell-type annotation or expert knowledge (**Fig. 2F**). Using in-house Xenium datasets, similarly we identified major tissues in the developing heart (**Supp. Fig. S2E**). Significant correlations were observed in the proportions of cell types between the matched tissue pairs when comparing the query data sets (Visium HD and Xenium-5K) with the reference datasets (**Fig. 2G**, **Supp. Fig. S2F**). The findings indicate that `TissueTypist` serves a robust and efficient cellular niche labelling tool, applicable to a diverse range of resolution datasets, including both sequenced and imaging-based spatial transcriptomics datasets. Using spatial omics datasets from other organs for training makes this workflow easily adaptable to different contexts.

### 2.3 Cellular circuits in the developing sinoatrial node

Pathway enrichment analysis of genes upregulated in SAN pacemaker cells (SANPCs) (compared with other cardiomyocytes) highlighted processes of the core pacemaking electrophysiological properties (**Fig. 3A, Fig. S3A**). We then constructed a gene regulatory network (GRN) targeting the identified pacemaker cell gene signatures using `SCENIC+`[19] and Multiome data. Centrality of the TF interactions highlighted well-described TFs for pacemaker cell development [20, 21], such as *ISL1*, *SHOX2*, *TBX3*, *TBX5*, *TBX18*, as hubs within the network. When we subsetted the GRN to the interactions targeting pacemaker-specific ion channel genes, including *HCN1*, *HCN4*, and *CACNA1D*, this also showed the high centrality of the well-described TFs (**Fig. S3B**). We then sought to explore foetal-specific properties of pacemaker cells by comparing the expression of the 675 upregulated pacemaker genes between foetal and adult pacemaker cells using our recent reference [22](**Fig. 3A**). We found 143 foetal-specific genes encompassing those involved in axon guidance processes, such as semaphorins and slit genes (*SEMA3A*, *SLIT2*, *SLIT3*)(**Fig. S3A**). We noted that these genes were highly expressed in the first-trimester hearts, with downregulation in the second-trimester and adult hearts (**Fig. 3B**). This suggests that pacemaker cells may play a significant role in the innervation of the developing heart. To understand the GRNs governing axon guidance gene expression in pacemaker cells, we used the `SCENIC+` output. The constructed network showed both TFs directly targeting these genes as well as high-level TFs, which may indirectly orchestrate expression (**Fig. 3C**). We noted that the ligand genes *SEMA3A*, *SLIT3*, *EFNB2*, and *CXCL12* were targeted by multiple TFs, suggesting that their regulation may involve cooperative TF interactions and/or redundancy mechanisms.

**Fig. 3:**
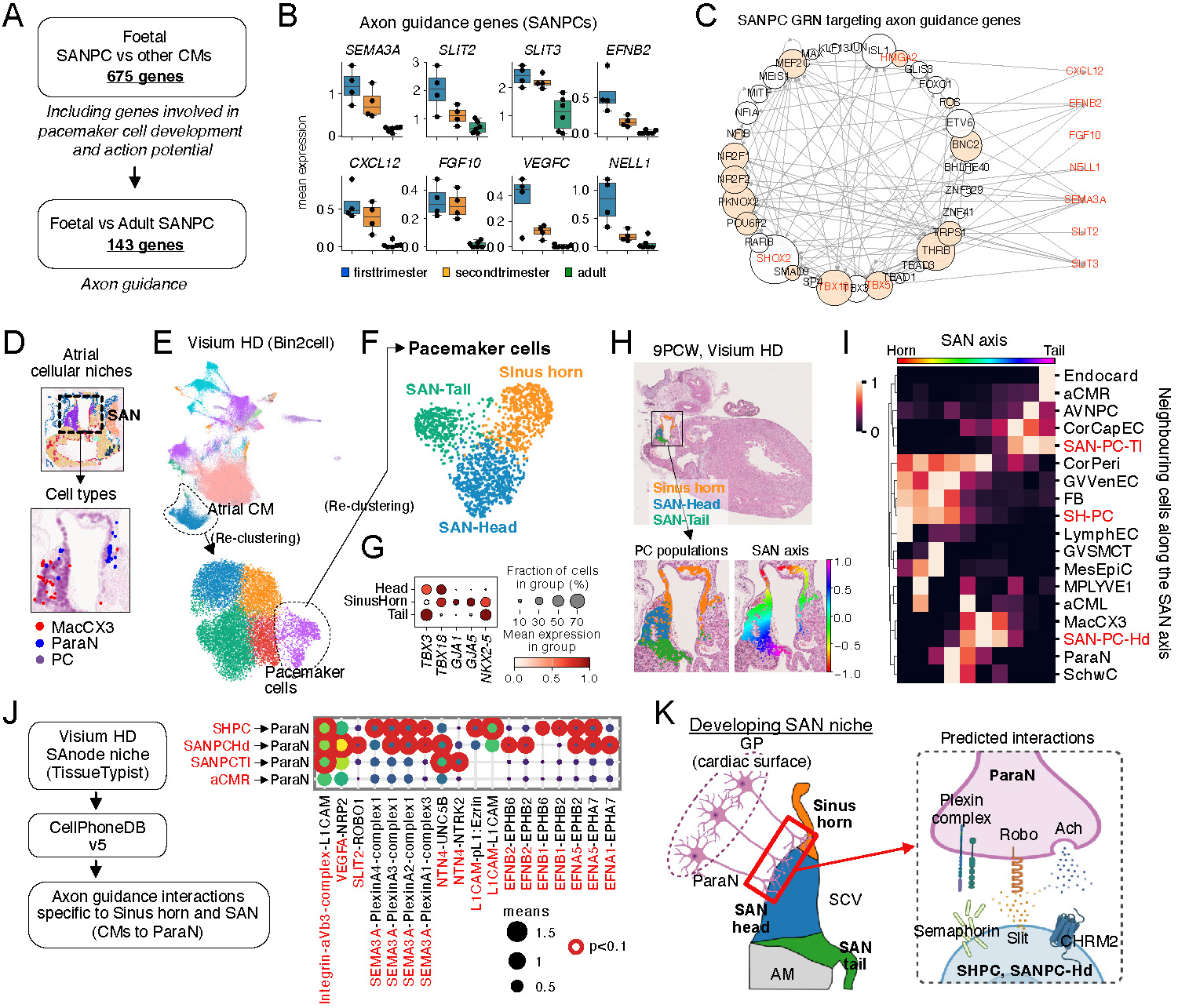
Cellular circuits in the developing sinoatrial node. **A:** Workflow of differentially expressed gene (DEG) analysis for SAN pacemaker cells. Pathways identified by gene-set enrichment analysis (Supp. Fig. 3A) are italicised. **B:** Expression of axon-guidance DEGs (foetal vs adult, adjusted p-value ¡ 0.05) in SAN pacemaker cells across foetal and adult [22] life stages. Dots show donor means; box-plot centre = median, box = 25–75 %, whiskers = min–max. **C:** Gene-regulatory network (GRN) targeting axon-guidance genes specifically expressed in SAN pacemaker cells. Transcription-factor (TF) nodes are sized by eigenvector centrality; salmon nodes denote TFs directly linked to target genes (TGs); red TGs are higher in foetal than adult pacemaker cells. **D:** Visium HD plot showing three cell types co-localise in the SAN. **E, F:** UMAP of Bin2cell transcriptomes (9 PCW). *SHOX2* -positive atrial cardiomyocytes extracted (E) and sub-clustered (F). **G:** Dot-plot of Bin2cell data showing marker expression for sinus-horn, SAN-Head and SAN-Tail pacemaker populations. **H:** Pacemaker populations and calculated SAN axis projected onto the Visium HD section. **I:** Normalised abundance of cell types neighbouring the SAN axis for SHPC, SANPC-Hd and SANPC-Tl. **J:** Inferred ligand–receptor interactions between atrial cardiomyocytes (pacemaker populations and right-atrial CMs) and parasympathetic neurons in the SAN niche; colour/size encode mean ligand–receptor expression; red border highlights cell-type-specific interactions. **K:** Schematic of the developing SAN niche and predicted ParaN–pacemaker interactions in the sinus horn and SAN-Head. **Abbreviations**: SAN – sinoatrial node; SANPC – SAN pacemaker cell; CM – cardiomyocyte; SHPC – sinus-horn pacemaker cell; ParaN – parasympathetic neuron; aCMR – right-atrial cardiomyocyte; CCS – cardiac conduction system; GP – ganglionated plexi; SCV – superior vena cava; AM – atrial myocardium; Ach – acetylcholine; CHRM2 – muscarinic acetylcholine receptor M2. Additional fine-grained cell-type abbreviations appear in Supplementary Table 6.

The SAN was profiled using Visium, Visium HD and Xenium. The SAN structure of the Visium HD data was localised between the epicardium and the developing superior vena cava (**Fig. 1E**). In both the Visium and Visium HD data, the SAN showed enrichment of pacemaker cells, parasympathetic neurons, and CX3CR1+ macrophages (**Fig. 2A**, **Fig. 3D**, **Fig. S3C-E**). We found that CX3CR1+ macrophages were predicted as a “microglia-like” population (**Supplementary Text**) and expressed microglia marker genes, including *CX3CR1*, *TREM2*, *P2RY12*, *ADGRG1* and *C3* (**Fig. S3F**). *CX3CL1*, the sole known ligand for CX3CR1, was specifically expressed by the parasympathetic neurons (**Fig. S3G**), suggesting CX3CR1^+^ macrophage are drawn to the SAN by parasympathetic neurons (the same interaction causes efficient migration of microglia to neurons in the CNS [23].

`Bin2cell` analysis of the Visium HD captured approximately 900 pacemaker cells. Subclustering these cells revealed three distinct subpopulations expressing markers corresponding to pacemaker cells of the sinus horn (SHPC), SAN head (SANPC-Hd), and SAN tail (SANPC-Tl)(**Fig. 3E-G**) [1, 20]. Although human foetal pacemaker cell subtypes have recently been profiled using snRNAseq [24] our data is the first to map them to their corresponding structures, confirming their annotation (**Fig. 3H**). Using the OrganAxis algorithm, we constructed a trans-SAN axis extending from the sinus horn to the SAN tail, revealing both cell types and gene expression along the axis (**Fig. 3H, I**; **Fig. S3H**). Parasympathetic neurons and CX3CR1+ microglia-like macrophages co-localise with the SHPC and SANPC-Hd, but not with SANPC-Tl (**Fig. 3I**). Several neuro-attractant signalling axes were specific to the SAN Head and sinus horn including SLIT-ROBO, Semaphorin 3A and Ephrin signalling, suggesting these pathways are important in drawing parasympathetic neurons to the SAN (**Fig. S3H**). Xenium data also demonstrated close localisation of SANPCs with parasympathetic neurons (**Fig. 1E**).

To further explore cellular crosstalk in the SAN during development, we performed cell-cell interaction analysis using the Visium HD (`CellPhoneDB`[25]), the workflow of which restricts ligand-receptor analysis based on spatially co-localised cell types and gene expression. In line with the observations noted previously, the inferred interactions indicated stronger connections between SANPC-Hd or SHPC and the parasympathetic neurons, especially through semaphorine and ephrin signalling, when compared to the interactions between working aCM and the parasympathetic neurons (**Fig. 3 J**). This suggests that neuronal cell recruitment signals originate from pacemaker cells. To validate that the expression profiles of each ligand and receptor align with the anticipated cell types, we conducted a `CellPhoneDB` analysis utilising snRNA-seq data. The results obtained were consistent with our predictions using the Visium HD data (**Fig. S3I**).

To elucidate the transition of pacemaker cell subpopulations throughout development, we identified these subpopulations in the single-nucleus RNA-seq data, leveraging the annotations of the Visium HD data and `CellTypist` (**Fig. S3J,K**). Our findings indicate that the SANPC-Hd population constitutes a substantial proportion of the early first trimester (**Fig. S3L**) while expressing higher levels of axon guidance genes along with SHPC (**Fig. S3M**), suggesting that these populations promote innervation during the early stages of development.

Recently, it has been suggested that para- and autocrine glutamatergic signalling may regulate firing rate in both working cardiomyocytes and pacemaker cells [22, 26]. Amongst the genes upregulated in adult compared to foetal pacemaker cells, we found several genes supporting glutamatergic signalling (*SLC1A3*, *SLC17A5*, *GRIA3*, *GRID2*). This supports a role for glutamatergic signalling in adult pacemaker cells and suggests that this capability is obtained after the second trimester (**Fig. S3N**).

Together, these findings suggest that the innervation of the SAN occurs in the region where SANPC-Hd cells are localised. SANPC-Hd cells express a series of genes promoting vagal innervation, particularly in the first trimester, suggesting the cells may play a significant role in this process. These data outline the developing SA nodal niche for the first time, showing evidence of crosstalk between colocalised pacemaker cells, parasympathetic neurons as well as microglia-like CX3CR1+ macrophages (**Fig. 3 K**).

### 2.4 Atrial cardiomyocyte maturation

Two transcriptionally distinct populations of working atrial cardiomyocytes were identified [10] (**Fig. 4A**) and Visium spatial mapping revealed that these populations were enriched in the left and right atria, respectively (aCMLs and aCMRs) (**Fig. S4A**). aCMLs were marked by expression of (*PITX2* and *PANCR*: PITX2-associated non-coding RNA) which have been highlighted in human and mouse left atrial tissue [27, 28], and are important in repressing the pacemaker gene programme in the left horn of the sinus venosus [21]. The *PITX2* -adjacent 4q25 locus is a well-known risk locus for atrial fibrillation [21]. aCMRs exhibit greater expression of genes involved in axon guidance and innervation (**Fig. S4B**), including Neurotrimin (*NTM*), SLIT2 and 3 and Semaphorin 5A (*SEMA5A*), all of which increased in expression in the second trimester (**Fig. 4D**). The cognate receptors were expressed by both sympathetic and parasympathetic neurons (**Fig. S4C**). This ligand expression difference in innervation-promoting signalling aligns with the greater density of autonomic innervation in the right atrium post-natally [16].

**Fig. 4:**
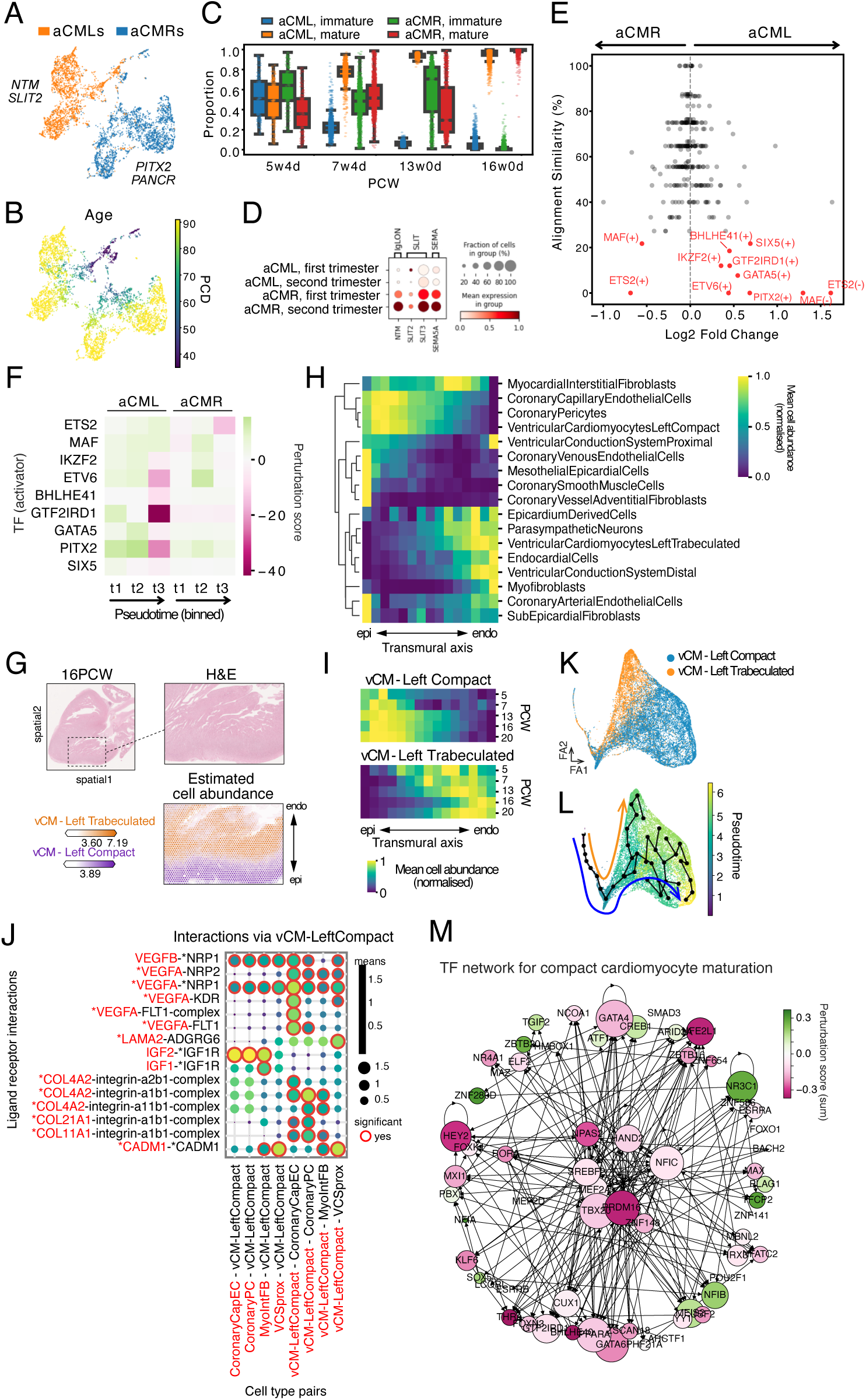
Atrial and ventricular cardiomyocyte development. **A, B:** UMAP embedding of atrial cardiomyocytes based on both gene expression and chromatin accessibility data (multiVI). Fine-grained cell type labels (aCMLs and aCMRs; left and right atrial cardiomyocytes)(A) or chronological age (PCD; post-conception days)(B) are projected on the UMAP. **C:** Boxplot showing proportions of immature and mature atrial cardiomyocytes over time in the atrial Visium spots spanning 5 to 16 PCW based on the cell type mapping (spot deconvolution) results. Four atrial carciomyocyte labels (immature and mature for each of aCMLs and aCMRs) were included in the analysis (**Fig. S4D**). For each spot, for each aCML or aCMR, the proportion of the mature or immature cell type was calculated using the total of immature and mature as the denominator (Methods). **D:** Dot plot showing expression of four axon guidance genes which are upregulated in aCMLs compared to aCMLs. Data is grouped by a concatenation of cell type and trimester. **E:** Scatterplot of genes2genes alignment results comparing regulon activity scores (obtained using SCENIC+) as features between aCML and aCMR trajectories. Low similarity (¡25%) regulons are highlighted in red. **F:** *In-silico* perturbation score (CellOracle) of the TFs highlighted using the genes2genes alignment (E). The perturbation score was obtained by using CellOracle and the atrial cardiomyocyte embedding (A). A negative score means that the TF perturbation will impede the atrial cardiomyocyte maturation trajectory. **G:** Abundance of compact and trabeculated cardiomyocytes at the left ventricle region of 16 PCW Visium section as estimated by cell2location. **H:** Cell type abundance transition across the transmural axis of Visium data. The colour scale shows the scaled value (min-max normalisation per cell type) of the mean cell abundance per transmural axis bin. **I:** Normalised cell abundance per spot along the left ventricular transmural axis and the chronological age. The transmural axis was binned and the mean of the values were calculated for each bin and age. **J:** Inferred cell-cell interactions between the left ventricular compact cardiomyocytes (vCM-LeftCompact) and the other cell types in the ventricular compact niche. The colour scale and dot size represent the mean expression levels of the interacting ligand-receptor partners. Cell type-specific interactions are delineated with a red border. Compact cardiomyocyte signature genes are highlighted with asterisks. **K, L:** Force-directed graph of both gene expression and chromatin accessibility data (MultiVI) for left ventricular compact cardiomyocytes. Fine-grained cell type labels (K) or calculated pseudotime (L) are projected on the graph. **M:** TF network targeting compact-specific and commonly upregulated signatures. GRN was constructed using SCENIC+, and the interactions for the TF network were selected as described in Methods. The size of the nodes represents the centrality score (eigenvector) in the TF network. The colour scale indicates the *in-silico* perturbation score obtained from CellOracle analysis (Methods). A negative score means that the TF perturbation will impede the trajectory (compact cardiomyocyte maturation). **Abbreviations**: aCML - left atrial cardiomyocytes; aCMR - right atrial cardiomyocytes.

To explore the maturation of atrial cardiomyocytes we constructed a `multiVI` embedding using both RNA and ATAC-seq data (**Fig. 4A**). Comparing this with chronological sample age revealed that aCML and aCMR profiles in younger hearts are relatively similar, and that their profiles diverge during development (**Fig. 4B**). Discretising the aCML and aCMR profiles into immature and mature versions using a 7 PCW cut-off (**Fig. S4D**), then mapping these into Visium from multiple developmental ages showed a clear trend where the immature states are gradually replaced by their mature counterparts, showing that spatial transcriptomic data support the age-dependent transition seen in the single-cell data (**Fig. 4C**).

To further examine this divergence, a pseudotemporal ordering was created based on the `multiVI` latent space, which correlated highly with chronological age (**Fig. S4E**). To compare differences between the trajectories we then used `genes2genes`, which identifies feature-level alignment between two trajectories [29]. We applied `genes2genes` to the `SCENIC+`-derived regulon activity scores. We reasoned that regulons with low similarity across the whole trajectory may potentially exert an early and sustained influence during development. This highlighted eleven highly mismatched regulons, nine of which had a higher mean activity in the aCML trajectory (**Fig. 4E**). Among these, the PITX2 regulon re-appeared, providing further confidence to our analysis. Interestingly, the activating and repressing activities of two regulons (ETS and MAF) showed reciprocal activity in the two trajectories. This suggests that these TFs may be bifunctional, promoting divergence towards two different cell identities. To predict the effect of TFs on the maturation of both aCML and aCMR, we performed an *in silico* TF perturbation analysis using `CellOracle` [30]. Knockout of the nine activator TFs identified through `genes2genes` analysis, including PITX2 and ETS2, revealed potential perturbation effects in both the aCML and aCMR lineages (**Fig. 4F**, **Fig. S4F**).

In summary, although the morphological atria have formed prior to our earliest sample (around post-conception day 28), we show that atrial cardiomyocytes continue to mature and the aCML and aCMR populations become more distinct with development.

### 2.5 Ventricular cardiomyocyte development

Single-cell data revealed compact and trabeculated populations of ventricular cardiomyocytes [10]. Spatial mapping validated these annotations, showing consistent overlap with their respective histological structures in the corresponding H&E images (**Fig. 4G**). To analyse the spatiotemporal transition of compact and trabeculated cardiomyocytes across the ventricular wall, we defined a transmural axis between the manually annotated epicardial and endocardial layers of the left ventricle using `OrganAxis` [11] (**Fig. S4G**). This provided a common spatial coordinate framework to compare features, such as genes or cell type abundances, across developmental time points and hearts of varying sizes (**Fig. 4H**).

During development, we found the abundance distribution of compact cardiomyocytes along this axis shifted from a tight distribution at the epicardium, to a more dispersed distribution closer towards the endocardium, while the inverse shift was observed for trabeculated cardiomyocytes (**Fig. 4I**). Additionally, nuclear density was seen to be highest in the older hearts particularly towards the epicardial boundary (**Fig. S4H**). Collectively, these findings suggest that the compact cardiomyocyte population expands in the second-trimester heart.

The compact cardiomyocytes showed a similar transmural distribution to coronary pericytes and capillary endothelial cells (which were constituents of the same ‘compact myocardium’ niche (**Fig. 4H**). Furthermore, these endothelial cells dispersed across the ventricular wall in tandem with the compact cardiomyocytes (**Fig. S4I**). Cell-cell interaction inference suggested that compact cardiomyocytes promote angiogenesis *via* VEGFs directed towards these coronary capillary endothelial cells (**Fig. 4J**). By contrast, trabeculated cardiomyocytes localised with centricular conduction system (VCS) cells, endocardial cells, and myofibroblasts at the endocardial boundary (**Fig. 4H**).

To clarify the molecular mechanisms involved in the maturation of compact and trabeculated cardiomyocytes, we utilised gene expression and chromatin accessibility data to create an integrated embedding of left ventricular cardiomyocytes (**Fig. 4K**). The integrated embedding showed the independent temporal transition for compact and trabeculated populations. Pseudotime was calculated by specifying the youngest tip principal point as the root, and the resulting pseudotime assignment correlated with the post-conception age of the samples (**Fig. 4L**). The genes associated with pseudotime in the compact and trabeculated cardiomyocyte trajectories were selected, and the common upregulated gene-set was enriched with genes involved in pathways related to vCM maturation, such as ‘response to calcium ion’ or ‘regulation of heart contraction’ (**Fig. S4J**). The compact-specific signature was enriched for processes involved in lipid transport and metabolism, such as *CD36*, *OSBPL2*, *PITPNC1* and *SREBF2*, suggesting a preparatory mechanism for a metabolic switch.

Within the GRNs targeting common and compact cardiomyocyte maturation, the centrality of the TF interactions showed several TFs as hubs within the network, such as PRDM16, TBX20, PPAR-*α*, GATA4 and GATA6 which have been implicated in cardiomyocyte maturation [31, 32] (**Fig. 4M**). Our analysis also highlighted TFs that are less well established within cardiomyocytes, including *SREBF2*, which encodes SREBP2. SREBP2 is a key regulator of sterol and fatty acid synthesis, a metabolic pathway that plays crucial roles in cardiomyocyte maturation [33]. To predict the effect of TFs in the maturation of compact cardiomyocytes, we performed an *in silico* TF perturbation analysis using `CellOracle` [30]. This highlighted the implication of several transcription factors, including HEY2, PRDM16, NFE2L1, and SREBF2 in the maturation of left compact vCMs (**Fig. S4K**).

Together, we employed multiomic analyses to investigate the spatiotemporal transition during ventricular cardiomyocyte development and to identify potential regulatory mechanisms responsible for the cardiomyocyte compaction.

### 2.6 Trisomy 21 disrupts compact myocardium formation through increased apoptosis

Congenital heart disease (CHD) encompasses a wide variety of cardiac structural defects affecting approximately 1% of live births [34]. Its aetiology is complex with both genetic and environmental drivers. Trisomy 21 (T21) is a strong risk factor for CHD, with a frequency of 40 to 50% amongst T21 live births and confers a particular risk for septal defects (interatrial, interventricular, or canal) [3]. A recent study showed impaired proliferation, compaction and mitochondrial respiration in cardiomyocytes using data from human and murine models, which were associated with heart septation defects [35]. However, the mechanisms underlying this phenotype remain incompletely understood.

To shed light on this we generated Multiome and Visium data from T21 hearts (aged 11-14 PCW). After quality control, there were 110,000 nuclei, which included a comprehensive set of cell types (annotated using a `CellTypist` [15] model trained on the euploid atlas). T21 nuclei were integrated with the age-matched euploid atlas using `scVI` (**Fig. 5A**, see the Methods section for a detailed description). The differential gene expression analysis for each cell type revealed a significant enrichment of differentially expressed genes (DEGs) on chromosome 21, as anticipated (**Fig. S5A**).

**Fig. 5:**
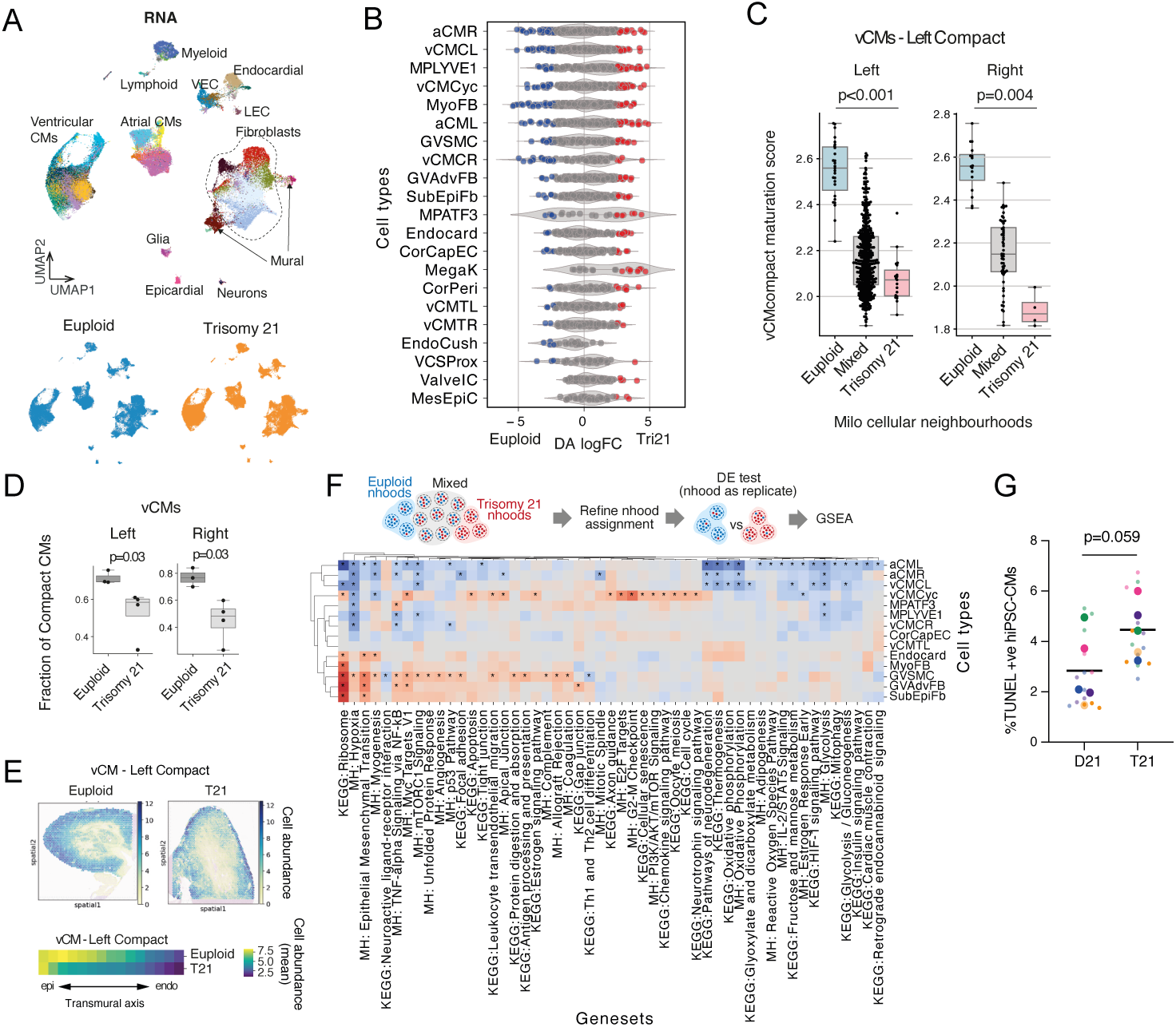
Trisomy 21 disrupts compact myocardium formation. **A:** UMAP embedding of gene expression data of the Trisomy 21 (11-14 PCW, 4 donors) and the age-matched euploid (3 donors) single-nucleus Multiome data. The fine-grained cell types are coloured, while the mid-grained cell types are labelled. **B:** Bee-swarm plot of log-fold change (x-axis) in cell abundance between euploid and T21 (scVI latent space). Cellular neighbourhoods overlapping the same cell type are grouped (y-axis). Neighbourhoods with significant differential abundance are coloured red or blue (Spatial FDR¡0.1). Cell types which have three or more significant neighbourhoods are selected and are ranked by the number of significant neighbourhoods. **C:** Box plot illustrating the calculated maturation scores for the compact cardiomyocyte cellular neighborhoods of the left ventricle. The assignment of neighborhoods has been refined to prevent the duplication of cells across multiple neighbourhoods. “Mixed” refers to the cellular neighbourhoods with no significant enrichment to either euploid or trisomy 21. **D:** Box plot showing the proportion of compact cardiomyocytes in the left and right ventricle. Each dot represents a donor (T21: 4 donors, euploid: 3 donors). **E:** Estimated abundance (cell2location) of compact cardiomyocytes at the left ventricle of 13 PCW euploid and Trisomy 21 Visium sections (top). Heatmaps show scaled cell abundances for the cell types across the transmural axis (bottom). **F:** Heatmap illustrating the results of Gene Set Enrichment Analysis (GSEA) derived from differential gene expression analysis utilising cellular neighbourhoods (Milo). The assignment of neighbourhoods has been refined to eliminate the duplication of cells across multiple neighbourhoods. The colour scale indicates the normalised enrichment score associated with Trisomy 21. “Mixed” refers to the cellular neighbourhoods with no significant enrichment to either euploid or trisomy 21. **G:** Dot plot illustrating the percentage of TUNEL-positive cardiomyocytes during the differentiation of human induced pluripotent stem cells (iPSCs) with Trisomy 21 (T21) compared to its isogenic control, which has disomy 21 (D21). Large dots represent biological replicates (different batches of differentiation), while small dots denote technical replicates. Statistical analysis was performed using the biological replicates. **Abbreviations**: DA logFC - differential abundance log2 fold change; vCM - ventricular cardiomyocyte; nhood - cellular neighbourhood; The abbreviations of the fine-grained cell types are listed in Supplementary Table 6.

As a measure of the difference between euploid cell types and their T21 counterparts, we compared their abundance at cellular neighbourhood level using `Milo` [36] (**Fig. 5B**). Cardiomyocytes exhibited high number of significantly enriched neighbourhoods, suggesting substantial transcriptional divergence in T21 (**Fig. 5B**). To evaluate the compaction and maturation of ventricular cardiomyocytes in our dataset, we scored each neighbourhood for the maturation and compaction gene signature from the euploid ventricular cardiomyocyte trajectory (**Fig. S4J**). The result suggests that ventricular cardiomyocytes in T21 hearts undergo reduced maturation and compaction, as indicated by lower score for maturation and compaction in T21-enriched neighbourhoods **Fig. 5C**. This is further supported by a decreased proportion of compact cardiomyocytes compared to euploid hearts (**Fig. 5D**). Consistent with these findings, the activity scores of GRNs involved in common and compact cardiomyocyte maturation (**Fig. 4M**) were significantly reduced in trisomy 21 compact cardiomyocytes when compared to euploid cells (**Fig. S5B**). Spatial transcriptomics analysis also showed a reduced estimated abundance of compact ventricular cardiomyocytes in T21 (**Fig. 5E**). To further explore why T21 hearts exhibit reduced compact ventricular cardiomyocytes, we performed cell-cell interaction analysis using `CellphoneDB`, comparing euploid and T21 cells. This highlighted a marked rewiring of Sempahorin–Plexin communication in T21 ventricles with upregulation of semaphorin ligand genes across multiple cardiac cell types in T21, including *SEMA3C* (**Fig. S5C**). This was accompanied by downregulation of *PLXNA4* and upregulation of *PLXNA2* in cardiomyocytes. Given prior work showing that SEMA3C–PLXNA2/4 signalling is required for myocardial compaction [9], this receptor–ligand mismatch may impair Semaphorin responsiveness which leads to reduced compact ventricular cardiomyocytes.

To better understand the differences between the two karyotypes, we performed differential gene expression analysis for each cell type within cellular neighbourhoods enriched for T21 and euploid cells (**Fig. 5F**). This revealed that pathways related to myogenesis, hypoxia response, and oxidative phosphorylation were disrupted in T21-enriched neighbourhoods. Additionally, apoptosis and cellular senescence pathways were upregulated in T21 cells, particularly within the cycling ventricular cardiomyocyte population. Endocardial cells, previously implicated as major contributors to the Down’s syndrome cardiac phenotype, exhibited alterations in epithelial-to-mesenchymal transition signatures.

To investigate conserved gene program perturbations between humans and mice, we generated scRNA-seq data from the hearts of E13.5 wild-type (WT) mice and Down syndrome mouse models (Dp1Tyb)[37] (**Fig. S6A**). By integrating the data using scVI (**Fig. S6B**) and performing cellular neighbourhood-based differential expressed gene analysis and geneset enrichment analysis, we showed shared enriched genesets between humans and mice, such as impaired oxidative phosphorylation and heightened activation of apoptosis pathways (**Fig. S6C**).

Survival and anti-apoptotic signalling closely associate with the proliferation, compaction, and maturation of cardiomyocytes [38]. Increased levels of apoptosis have been linked to CHDs and the impairment of myocardial compaction[39]. To assess the impact of T21 on apoptosis sensitivity, we differentiated human T21-induced pluripotent stem cells (T21-iPSCs) and their isogenic counterparts into cardiomyocytes. Using TUNEL staining we evaluated the proportion of apoptotic cardiomyocytes in these cultures and found an increased proportion of apoptotic cardiomyocytes within T21 cultures (**Fig. 5G**), suggesting that indeed T21 promotes apoptosis in cardiomyocytes.

These results demonstrate that trisomy 21 is associated with increased apoptosis of cardiomyocytes, potentially leading to reduced compaction of ventricular cardiomyocytes in T21 hearts compared to euploid hearts.

## 3 Discussion

Here we present the most comprehensive multimodal atlas of human cardiogenesis to date. Combining gene expression, chromatin accessibility and spatial transcriptomics, we shed light on dynamic processes occurring in human cardiogenesis. Compared to previous multiomics studies targeting developing human hearts [7–9], this study presents the most extensive human-heart multiomics atlas to date, spanning the first and second trimesters. In particular, leveraging high-resolution spatial transcriptomics with whole-transcriptome profiling, our analysis captures detailed cellular architecture, including the microenvironments of the SAN, at an unprecedented scale. We also introduce `TissueTypist`, a novel machine-learning tool that efficiently predicts cellular niches across spatial datasets of different resolutions, leveraging both gene expression and spatial context. Finally, comparative analysis of euploid and T21 samples reveals decrease of ventricular compact cardiomyocytes in the T21 hearts, implicating apoptosis-related mechanisms in congenital heart disease.

`TissueTypist` offers several advantages: it is applicable across spatial scales and platforms; it does not require pre-existing cell segmentation or manual annotation to operate; and it is fast and computationally efficient, enabling rapid analysis. The primary limitation is that the bundled model is trained on foetal heart tissues, though users can readily train custom models for other contexts. Additionally, as a classifier, `TissueTypist` does not enable discovery of novel tissue types. However, low-confidence predictions may indicate unseen structures, providing a basis for further exploration.

Using a combination of spatial transcriptomic platforms, we show that foetal pacemaker cells exhibit higher expression of axon guidance genes compared to their adult counterparts and colocalise with parasympathetic, but not sympathetic, neurons. These findings suggest that the first trimester is a critical period for parasympathetic innervation of the SAN. This period also aligns with the first trimester heart rate decrease which may be partly driven by the suppressive influence of this innervation [40]. It is consistent with previous animal studies which suggests parasympathetic innervation begins in the first trimester and precedes sympathetic innervation [41]. Using our paired single-nucleus RNA-seq and ATAC-seq data, we constructed enhancer-mediated GRNs governing axon guidance gene expression. This analysis highlighted the central role of canonical transcription factors—such as TBX3, TBX18, SHOX2, and ISL1—that are established drivers of pacemaker cell identity in mouse [42]. Their prominent involvement in regulating axon guidance genes in human pacemaker cells suggests that axon guidance is an integral component of pacemaker cell biology [43, 44]. Leveraging single-cell–resolved spatial transcriptomics (Visium HD and Xenium), we generated the first high-resolution map of the developing human SAN and its adjoining structures. Analysis along a trans-SAN axis revealed a pronounced enrichment of parasympathetic fibres in the sinus horn and, most strikingly, within the SAN head. Notably, the head domain is where the core pacemaker cardiomyocytes reside, as shown in both developmental and adult studies [45]. The concentration of parasympathetic input and axon guidance gene expression at this distal pacemaker compartment—rather than at the SAN-atrial border—implies that autonomic modulation acts directly on the dominant pacemaker region, offering an anatomical basis for fine-tuning of heart-rate control.

During the first and second trimesters, left- and right-atrial cardiomyocytes diverge along distinct transcriptional and epigenetic trajectories. Multi-omic integration shows that the left-atrial lineage (aCML) progressively up-regulates a PITX2-centred regulon, repressing the pacemaker programme while the right-atrial lineage (aCMR) acquires a strong axon-guidance and innervation signature (e.g. NTM, SLIT2/3, SEMA5A), consistent with the higher density of autonomic fibres in the postnatal right atrium [16, 27, 28]. Feature-level trajectory alignment highlighted nine regulons—including PITX2 and bifunctional ETS/MAF modules—that are already mismatched at the earliest time-points in our data (4 PCW), implying that left-right atrial identity is established by an early, sustained transcriptional landscape.

In the ventricle, spatially resolved single-cell data reveal a progressive expansion of compact cardiomyocytes from the epicardium towards the endocardium during the first and second trimesters, mirrored by a reciprocal retreat of trabeculated cardiomyocytes. Compact cardiomyocytes co-localise with coronary capillary endothelial cells and express VEGFA, suggesting a feed-forward coupling between myocardial thickening and angiogenesis. Pseudotemporal analysis uncovered a shared maturation module enriched for calcium-handling genes, but compact-specific signatures are dominated by lipid-metabolic regulators (CD36, SREBF2, OSBPL2), indicating that possibly compact myocardium undergoes a metabolic switch earlier. GRN inference and *in silico* perturbations nominate HEY2, PRDM16, NFE2L1 and SREBF2 as central drivers of this maturation programme.

Trisomy 21 (T21) is a major risk factor for congenital heart disease, and is particularly associated to defects of the muscular septum [46]. We present the first single-cell multiome and spatial transcriptomic data of cardiogenesis in T21. We show T21 leads to a pronounced depletion of compact cardiomyocytes, coronary endothelial cells and pericytes. Cellular neighbourhood-level gene-set enrichment reveals coordinated down-regulation of myogenic, oxidative-phosphorylation and hypoxia-response programmes, and reciprocal up-regulation of unfolded-protein-response and apoptosis pathways. *In vitro* validation confirms heightened TUNEL positivity, indicating that an intrinsic increase in apoptosis accompanies, and possibly is responsible for the reduction in ventricular compact cardiomyocytes. Mechanistically, our data complements and extends findings from the Dp1Tyb mouse model, in which an extra DYRK1A copy limits cardiomyocyte proliferation, mitochondrial function and septation [35]. Our data further suggests that elevated apoptosis is a factor hindering ventricular compaction, which may contribute to some of the septum defects typically seen in T21 individuals. Indeed it has been previously shown that ventricular septal defects can be associated with abnormalities in myocardial compaction[47–49].

Differential abundance analysis of human single-cell multiome data from T21 and euploid hearts revealed that cardiomyocytes appeared more affected than endocardial (endothelial) and endocardial cushion (mesenchymal) cells (**Fig. 5B**, **Fig. S6B**), the latter two being cell types previously implicated as major contributors to the Down syndrome cardiac phenotype. This observation aligns with prior reports indicating that Trisomy 21 disrupts multiple cardiac lineages, which together contribute to structural malformations [35]. In terms of qualitative differences, neighborhood-based differential gene expression analysis between euploid and T21 cell types identified alterations in epithelial-to-mesenchymal transition (EMT) signatures within endocardial cells (**Fig. 5F**); notably, EMT appeared increased in T21. The functional significance of this upregulation, however, remains to be determined. A key limitation is that atrial or ventricular septal defects, hallmark lesions in Down syndrome hearts, were not identified in our samples in histological sections. Because confirmation of such defects generally requires three-dimensional imaging of the heart, which was not undertaken, their presence cannot be definitively excluded. Future studies incorporating volumetric imaging will, therefore, be needed to capture the full spectrum of structural anomalies.

Overall, we present an integrative analysis of a high-resolution multi-omic map of the developing human heart during the first and second trimesters. This resource and the insights presented here are anticipated to broaden our understanding of human heart development with a range of implications in disease modelling and regenerative medicine, as exemplified by our insight into how T21 contributes to CHD.

## 4 Methods

### 4.1 Human embryonic and foetal heart tissues

All tissue samples were obtained after Research Ethics Committee approval and written informed consent from mothers. The embryonic and foetal heart samples corresponding to the donor IDs BRC2251, BRC2253, BRC2256, BRC2260, BRC2262, BRC2263, C82, C83, C85, C86, C87, C92, C94, C97, C98, C99, C104, and C194 were provided from terminations of pregnancy from Cambridge University Hospitals NHS Foundation Trust under permission from NHS Research Ethical Committee (96/085). The donor IDs Hst32, Hst33, Hst36, Hst39, Hst40, Hst41, Hst42, Hst44, Hst45, and Hst48 were provided from the MRC/Wellcome Trust Human Developmental Biology Resource (University College London (UCL) site REC reference: 18/LO/0822; www.hdbr.org. Registered project title: A multi-omics analysis of fetal cardiac development and the impact of Trisomy 21, Registered project number: 200556 and 200747). Donor IDs and that information are listed in Supp. Table 1.

### 4.2 Sample collection and processing

The processing of samples with donor IDs of BRC2251,BRC2253, BRC2256, BRC2260, BRC2262 and BRC2263 were described previously^71^. Briefly, these samples were stored at 4 °C overnight in Hibernate-E medium (ThermoFisher Scientific). The next day, the apex and the base of the heart were dissected and separately dissociated using 6.6 mg/mL Bacillus licheniformis protease (Merck), 5 mM CaCl2 (Merck), and 20 U/mL DNase I (NEB). The mixture was triturated on ice for 20 seconds every 5 minutes until clumps of tissue were no longer visible. The digestion was stopped with ice-cold 10% fetal bovine serum (FBS, ThermoFisher Scientific) in phosphate-buffered saline (PBS, ThermoFisher Scientific). Cells were then washed with 10% FBS, resuspended in 1 mL PBS and viability assessed using Trypan blue. Cells were submitted for 10x library preparation with v3.0 chemistry for 3’ single-cell sequencing on a NovaSeq 6000 (Illumina) at the Cancer Research UK Cambridge Institute.

For the processing of donors corresponding to the donor IDs C86, C94, C97 and C99, samples were stored at 4 °C overnight in HypoThermosol preservation solution (Merck). Tissue was first minced in a tissue culture dish using scalpel. Minced tissue was digested with type IV collagenase (final concentration of 3 mg/mL; Worthington) in RPMI (Sigma-Aldrich) supplemented with 10% fetal bovine serum (FBS; Gibco), at 37°C for 30 min with intermittent agitation. Digested tissue was then passed through a 100-µm cell strainer, and cells were pelleted by centrifugation at 500g for 5 min at 4°C. Cells were then resuspended in 5 ml of red blood cell lysis buffer (eBioscience) and left for 5 min at room temperature. It was then topped up with a flow buffer (PBS containing 2% (v/v) FBS and 2 mM EDTA) to 45 ml and pelleted by centrifugation at 500g for 5 min at 4°C. The resuspended cell solution was filtered through a 70-*µ*m cell strainer (Corning), and live cells were manually counted by Trypan blue exclusion. Dissociated cells were first incubated with 5 *µ*L of FcR blocker for 5 min at room temperature and stained with anti-CD45 antibody (BV785 anti-human CD45 antibody, BioLegend, 304048) and DAPI (Sigma-Aldrich, D9542) prior to sorting. DAPI was used at a final concentration of 2.8 *µ*M, and all antibody solutions were used at a final concentration of 5 µl per 100 *µ*l cell suspensions containing fewer than 5 million cells. DAPI-CD45+ and DAPI-CD45- populations were sorted by FACS using MA900 Multi-Application Cell Sorter (Sony) and its proprietary software (Cell Sorter Software v3.1.1). Sorted cells were loaded on the Chromium Controller (10x Genomics) with a targeted cell recovery of 5,000–10,000 per reaction. Single-cell cDNA synthesis, amplification, gene expression library was generated according to the manufacturer’s instructions of the Chromium Next GEM Single Cell 5’ Kit v2 (10x Genomics). Libraries were sequenced using NovaSeq 6000 (Illumina) at Wellcome Sanger Institute with a minimum depth of 20,000–30,000 read pairs per cell.

Samples used for single nuclei isolation and spatial transcriptomics were flash-frozen (unembedded) or frozen in OCT and stored at -80 °C, or formalin-fixed and subsequently embedded in paraffin blocks. All tissues were stored and transported on ice at all times until freezing or tissue dissociation to minimise any transcriptional degradation. Single nuclei were obtained from flash-frozen tissues using sectioning and mechanical homogenization as previously described. 5-10 mm thickness frozen tissues were first sectioned with cryostat in a 50 µm thickness section. All sections from each sample were homogenised using a 7 ml glass Dounce tissue grinder set (Merck) with 8-10 strokes of a tight pestle (B) in homogenization buffer (250 mM sucrose, 25 mM KCl, 5 mM MgCl2, 10 mM Tris-HCl, 1 mM dithiothreitol (DTT), 1× protease inhibitor, 0.4 U *µ*L-1 RNaseIn, 0.2 U *µ*L-1 SUPERaseIn, 0.05% Triton X-100 in nuclease-free water). Homogenate was filtered through a 40-µm cell strainer (Corning). After centrifugation (500g, 5 min, 4 °C) the supernatant was removed and the pellet was resuspended in storage buffer (1× PBS, 4% bovine serum albumin (BSA), 0.2U *µ*l-1 Protector RNa-seIn). Nuclei were stained with 7-AAD Viability Staining Solution (BioLegend) and filtered through 20-µm cell strainer (CellTrics). Positive single nuclei were purified by fluorescent activated cell sorting (FACS) using MA900 Multi-Application Cell Sorter (Sony) and its proprietary software (Cell Sorter Software v3.1.1). Nuclei purification and integrity were verified under a microscope, and nuclei were manually counted by Trypan blue exclusion. Nuclei suspension was adjusted to 1000-3,000 nuclei per microlitre and loaded on the Chromium Controller (10x Genomics) with a targeted nuclei recovery of 5,000-10,000 per reaction. 3’ gene expression libraries and ATAC libraries were prepared according to the manufacturer’s instructions of the Chromium Multiome ATAC+Gene Expression Kits (10x Genomics). Libraries were sequenced using NovaSeq 6000 (Illumina) at Wellcome Sanger Institute with a minimum depth of 20,000–30,000 read pairs per nucleus.

### 4.3 Spatial transcriptomics processing

Fresh-frozen (FF) samples: Fresh samples were frozen and embedded in optimal cutting temperature medium (OCT) using a dry ice-cooled bath of isopentane at -45°C. OCT-embedded samples were sectioned using a cryostat (Leica CX3050S) and were cut at 10 *µ*m. Formalin-fixed paraffin-embedded (FFPE) samples: Fresh samples were fixed in ¿5 times their volume of 4% v/v formalin at ambient temperature for 24 hours before processing to paraffin on a Tissue-Tek Vacuum Infiltration Processor 5 (Sakura Finetek). FFPE blocks were sectioned at 5 *µ*m using a microtome (Leica RM2125RT). RNA integrity number (fresh-frozen samples) or DV200 (formalin-fixed) was obtained using Agilent 2100 Bioanalyzer. CytAssist FFPE Visium Spatial Gene Expression (10x genomics) was performed following the manufacturer’s protocol. For FF samples, the Tissue Optimization protocol from 10x Genomics was performed to obtain a permeabilisation time of 45 min, and the FF Visium Spatial Gene Expression experiment was performed as per the manufacturer’s protocol (10x Genomics). H&E stained Visium Gene Expression slides were imaged at 40× on Hamamatsu NanoZoomer S60. After transcript capture, Visium Library Preparation Protocol from 10x Genomics was performed. Eight cDNA libraries were diluted and pooled to a final concentration of 2.25 nM (200 *µ*l volume) and sequenced on 2× SP flow cells of Illumina NovaSeq 6000. For Visium HD, the tissue section was prepared as same as the standard Visium samples. H&E staining and imaging were performed following the Visium HD FFPE Tissue Preparation Handbook (CG000684). Samples were then processed and sequenced following the Visium HD Spatial Gene Expression Reagent Kits User Guide (CG000685). The Xenium In Situ Gene Expression with Morphology-based Cell Segmentation Staining User Guide (CG000749) with the Xenium Prime 5K Human Pan Tissue & Pathways Panel (5001 genes).

### 4.4 Read mapping

After sequencing, samples were demultiplexed and stored as CRAM files. Each sample of single-cell data was mapped to the human reference genome (GRCh38-2020-A) provided by 10x Genomics using the CellRanger software (CellRanger v.6.0.2 or v.6.1.0, or CellRanger ARC v.2.0.0) with default parameters. Each sample of Visium was mapped to the human reference genome (OCT: GRCh38-3.0.0, FFPE: GRCh38-2020-A) the SpaceRanger software (OCT: v.1.1.0, FFPE: v.2.0.0, Visium HD: v.3.0.1) with default parameters. For Visium samples, SpaceRanger was also used to align paired histology images with mRNA capture spot positions in the Visium slides. Part of the single-cell samples were mixed with different donors after the nuclei isolation for cost-efficient experimental design (Supp Table 2) and computationally demultiplexed (Soupercell, v.2.0)^72^ based on genetic variation between the donors.

### 4.5 Preprocessing and quality control

For the single-cell transcriptome data, the CellBender algorithm (remove-background)(v.0.2)[50] was applied to remove ambient and background RNA from each count matrix produced using the CellRanger pipeline. Downstream analysis was performed using the Scanpy package. Doublets with a score of ¿0.15 were removed using Scrublet[51]. We performed quality control and filtering based on the following settings: number of genes¿500, total counts¿1000, % mitochondrial¡5 (nuclei), % mitochondrial¡20 (cell), % ribosomal¡5 (nuclei), % ribosomal¡20 (cell), red blood cell score¡1 (nuclei), red blood cell score¡2 (cell). For the multiome ATAC data, the data processed using CellRanger ARC were further analysed using ArchR (v.1.0.2)[52]. Quality control was performed, considering, among other factors, transcription start site enrichment, nucleosomal banding patterns, the number and fraction of fragments in peaks, reads falling into ENCODE blacklist regions as well as doublet scores computed by ArchR. For high-quality cells, reads were mapped to 500-bp bins across the reference genome (hg38)(TileMatrix). Gene scores based on chromatin accessibility around genes were computed from TileMatrix using the createArrowFiles function to check their consistency with measured expression values. Before peak calling, pseudobulk replicates were generated (addGroupCoverages) for each fine-grained cell state annotated using the paired gene expression data. Peak calling (501 bp fixed-width peaks) was performed for each cell state, and the peak sets were merged to obtain a unified peak set (addReproduciblePeakSet). A cell-by-peak count matrix was obtained using the addPeakMatrix function. For the visium data, the histological tissue annotation was performed based on the H&E image, and the visium spots on the irrelevant regions (‘Out of tissue’ and ‘Cavity’) were removed. Spots of each sample were further filtered for more than 500 UMI (OCT) or 5000 UMI (FFPE) counts and 300 (OCT) or 1000 (FFPE) genes. For Visium HD data, Bin2cell[**?**] (v.0.3.3), which is a tool to reconstruct cells from the 2 *µ*m bin data by utilising H&E image and gene expression data, was performed to identify each single cell. Initially, the data was pre-processed by removing uninformative genes and bins (genes expressed in fewer than 3 cells and bins with no counts). For nucleus segmentation using H&E image, the input image resolution was controlled by the “microns per pixel” (mpp) parameter (mpp=0.3). We applied the b2c.destripe function with quantile=0.99 to correct the striped appearance. Nucleus segmentation was performed using StarDist, an efficient segmentation algorithm with a pretrained HE model: b2c.stardist with prob thresh=0.05. To capture transcripts in the cytoplasms around the identified nuclei, the b2c.expand labels function was used (max bin distance=2). To detect cells with missing nuclei in the H&E image, Bin2cell performed segmentation on a representation of total expression per bin. The total counts (destriped counts) per bin were converted into an image using the b2c.grid image function (mpp=0.3, sigma=5). Segmentation was then performed on the converted image with the pretrained StarDist fluorescence model: b2c.stardist with prob thresh=0.05 and nms thresh=0.5. The cell segmentations obtained from the H&E image and gene expression data were merged using the b2c.salvage secondary labels function. Counts of bins assigned to each cell were grouped using the b2c.bin to cell function to generate a cell-by-gene matrix. Using the reconstructed cells, we conducted quality control and filtered the cells according to the following criteria: number of genes ¿ 30, number of bins ¿ 3, and number of bins ¡ 60. The thresholds were chosen arbitrarily based on the distributions to exclude outliers. This resulted in a dataset containing 129,803 cells. For Xenium data, multimodal cell segmentation algorithm was applied to generate cell-by-gene matrix (Xenium instrument software version: 3.1.3.1, Analysis version: xenium-3.1.0.4). Quality control was performed, and cells with fewer than 50 genes were removed. This resulted in a dataset with 362,277 cells.

### 4.6 Spatial mapping of the fine-grained cell types

For the low-resolution Visium data, we used cell2location (v.0.1) method[53] to spatially map the developing heart fine-grained cell types defined using single-cell transcriptomics data[10] in the Visium data. In brief, we first estimated reference signatures of cell types using sc/snRNA-seq data and a negative binomial regression model provided in the cell2ocation package. The inferred reference cell type signatures were used for cell2location cell-type mapping that estimates the abundance of each cell type in each Visium spot by decomposing spot mRNA counts. The mapping was performed per group of sections of each time point (4, 5, 7, 13, 16, and 20 PCW). The H&E images of the Visium slides were used to determine the average number of nuclei per Visium spot (n = 20) in the tissue and used as a hyperparameter in the cell2location pipeline. For the Trisomy 21 (T21) Visium data, we used snRNA-seq data from T21 hearts as the reference for cell2location mapping, following the same workflow. For the immature versus mature atrial cardiomyocyte analysis, a temporary (i.e. used only for this analysis) annotation was made where the scFates milestones prior to the divergence of the principal graph were labelled as AtrialCardiomyocytesImmature (all other fine grain annotations remained) (Supp. Fig. 6C). These signatures were mapped in the Visium data using cell2location. The per spot cell type abundances were extracted for spots with the structural annotation ‘atria’. For each of the 4 labels (immature/- mature for each of L/RaCM) a proportion was calculated: the abundance of that label divided by the combined total of the immature and mature abundances for the corresponding cell type. For instance proportion_LaCMimmatur_e = abundance_LaCMimmature_ / (abundance_LaCMimmature_ + abundance_LaCMmature_). This enables a comparison between the mature and immature versions of the same cell type. For both Visium HD and Xenium datasets, cell-by-gene expression matrices were used to perform cell type annotation. Coarse-grained and mid-grained cell types were initially assigned using a label transfer algorithm (CellTypist[15]), with reference to single-cell and singlenucleus RNA-seq data. For each mid-grained cell type, highly variable genes were identified, followed by principal component analysis (PCA), batch correction using Harmony[54], construction of a neighbourhood graph, and clustering with the Leiden algorithm. Fine-grained cell types were subsequently annotated based on these clusters through marker gene expression and manual curation. Clusters exhibiting expression of marker genes from multiple cell types were labelled as “unclassified” or “doublet” and excluded from downstream analyses.

### 4.7 Single-cell data integration

After pre-processing and removal of low-quality cells and nuclei, single-cell transcriptomics data were integrated using scVI[55, 56] accounting for categorical covariates of donor, cell or nuclei, and kit 10x, as well as continuous covariates of total counts, %mito, and %ribo. Neighbourhood identification, high-resolution leiden clustering, and further dimensional reduction using UMAP were performed based on the scVI latent space. Cells and nuclei of extra-cardiac origin were removed based on marker gene expressions: lung epithelial cells (IGFBP2, EPCAM, SOX2, and NKX2-1); hepatocytes (ALB, APOA1, TTR); hepatic stellate cells (CRHBP, FCN3, OIT3); parathyroid cells (PTH, MAFB, GATA3); and further erythrocytes (FTH1, HBA1, HBA2, HBB). The remaining data were integrated using scVI with the same variables, followed by the downstream processing. To annotate cell types, cell barcodes comprising major clusters were subsampled from the initial raw count matrix for sub-clustering analysis and processed separately through the same downstream analysis as described. Subclusters were annotated based on immunohistochemical validation and literature evidence. For ATAC data, post-QC data were integrated using cell-by-peak count matrix peakVI(Ashuach et al. 2022) accounting for batch variation (donor). Neighbourhood identification and dimensional reduction using UMAP were performed based on the peakVI latent space.

### 4.8 Epigenetic stability

Multiome data of each mid-grained cell type was separately integrated for the gene expression (GEX) or chromatin accessibility (ATAC) data using scVI or peakVI, accounting for batch variation (donor and sampled region). Overlapping Milo neighbourhoods were identified using the calculated neighbourhood graph based on the scVI (n neighbors=15) or peakVI (n neighbors=100) latent spaces (*milo.make nhoods*). For each gene expression neighbourhood, the entropy of the distribution of corresponding nuclei across ATAC neighbourhoods was calculated (*scipy.stats.entropy*). The effects of GEX neighbourhood size (number of nuclei per neighbourhood) and variance of nucleus neighbours in the scVI latent space on the entropy score were regressed out using a multiple linear regression model (*sklearn.linear model.LinearRegression*). The corrected entropy score indicates the extent of diffusion of a group of transcriptionally similar nuclei in the ATAC space, which can be interpreted as the epigenetic stability of the group of nuclei.

### 4.9 Visium data integration and tissue type annotation

Both low-resolution visium OCT and FFPE data (4, 5, 7, 13, and 16 PCW) were integrated using spot-by-cell type estimated abundance (cell2location) matrix and BBKNN^76^ accounting for batch variation (section). Batch-balanced neighbour graph was used for leiden clustering and UMAP visualisation. To annotate tissue niches, major tissues were first annotated based on the clustering result and the estimated cell type abundances. Each major cluster was subsampled and subjected to the downstream workflow as described to annotate finer-grained tissues. Some annotations were fully (AVN, lymph node, and epicardium) or partly (AV ring, AV valves, aortic valve, valve apparatus, coronary artery, and great vessels) defined based on the histological tissue structures manually annotated using the paired H&E images.

### 4.10 TissueTypist

All the functions and documentations are available at the github repository (https://github.com/Teichlab/TissueTypist).

#### 4.10.1 Model training

- Reference dataset: the integrated Visium OCT data from four hearts, comprising twenty sections and eighteen cellular niches, were used.
- Feature selection: key tissue genes were identified using the highly variable gene detection method (scanpy.pp.highly variable genes) and through the analysis of differentially expressed genes across niches. This approach yielded a total of 2603 genes selected for model training.
- Gene expression of neighbouring spots: for each spot, six neighbouring spots were identified, and the maximum expression value among these neighbours was recorded for each gene. This process resulted in 2603 gene features derived from the individual spots and an identical number of “neighbouring” gene features.
- Distance to tissue edge: the tissue edge for each section was determined by calculating the number of direct neighbours per spot and applying a specified threshold. Spots situated on technical edges were flagged and excluded from further analysis. A k-nearest neighbours (kNN) graph was constructed using the identified tissue edge spots, followed by the computation of nearest-edge distances (Euclidean distance) for non-edge spots. Spots on the edges were assigned a distance of zero. To ensure consistency across sections, min-max normalisation and a log1p transformation were applied for consistency across sections.
- A logistic regression model (sklearn.linear model.LogisticRegression) was trained using the aforementioned data, incorporating varying weights for the gene expression profiles of neighbouring regions and the calculated distances to the tissue edge.
- To evaluate the performance of the tissue classification model, we performed stratified k-fold cross-validation (with k = 5) using the annotated reference dataset. In each fold, the dataset was partitioned into training (80%) and validation (20%) sets while preserving the proportion of tissue labels. The model was trained on the training subset and evaluated on the held-out validation subset. Overall classification accuracy was computed as the proportion of correctly predicted labels across all validation samples in each fold. To assess label-specific performance, we calculated the F1 score for each tissue label in each fold, using the harmonic mean of precision and recall.
- For predicting tissues of an imaging-based spatial transcriptomics data utilising a targeted gene panel (Xenium or MERFISH), the reference Visium dataset was processed after subsetting the genes included in the panel, and a model was subsequently trained.

#### 4.10.2 Prediction

- Generation of pseudo-bulk gene expression data: for single-cell-level spatial transcriptomics data, pseudo-bulk gene expression data was generated using a sliding window approach. Specifically, the data were segmented into adjacent windows to match the size of the low-resolution visium data, utilising the ‘squidpy.tl.sliding window’ function (reference for squidpy). For each defined window, gene expression data were aggregated through summation. The resulting aggregated counts were then normalized and log-transformed to prepare them for further analysis.
- For Visium HD data, 8 *µ*m-bin data was used for the generation of pseudo-bulk data and the prediction. The predicted labels were transferred to bin2cell object based on their spatial coordinates.
- Query data (either low-resolution Visium data or aggregated data) were processed in a manner identical to that of the reference data during model training. The models, trained with the corresponding gene panel, were subsequently employed for predictions.
- To assess whether the predicted tissue labels captured expected cellular composition, we calculated Pearson correlations between cell-type proportions per spatial niche in the query dataset (Visium HD or Xenium-5K) and corresponding proportions in the annotated reference Visium datasets.

### 4.11 Trajectory analysis

We first obtained a latent space based on both the RNA-seq and ATAC-seq data using MultiVI[57]. The latent space was used for obtaining two-dimensional embeddings and a series of trajectory analyses using scFates[58]. For left ventricular cardiomyocytes, multiome data of compact and trabeculated cardiomyocytes were integrated with accounting for categorical covariates of donor and sampled region, as well as continuous covariates of total counts, %mito, %ribo, and cell cycle scores (S and G2/M phase). Multiscale diffusion space was obtained using the MultiVI latent space and the Palantir algorithm[59]. The neighbourhood graph was obtained using the Palantir diffusion space and was used for force-directed graph embedding. The Palantir diffusion space was also used for obtaining a principal graph with SimplePPT algorithm. Pseudotime calculation and the assignment of milestones and segments were performed by setting the youngest milestones as the root. The genes whose expressions were associated and upregulated along the pseudotime were selected either by subsampling compact or trabeculated populations or by using all nuclei. For atrial cardiomyocytes, multiome data of left and right cardiomyocytes were integrated with multiVI accounting for the same covariates as before. To avoid an over-representation of either cell type, data where the whole heart had been processed was used. The downstream steps were performed similarly to the analysis above. Two trajectories, consisting of milestones leading from the root to terminal milestones in the left and right atrial cardiomyocyte clusters, respectively, were defined. Association of gene expression with pseudotime was tested along each of these two trajectories independently, as well as across the whole embedding. For monocyte and macrophages, both the cells and nuclei samples with region==’whole heart’ were integrated with accounting for categorical covariates of donor, cell or nuclei, and kit 10x, as well as continuous covariates of total counts, %mito, %ribo, and cell cycle scores (S and G2/M phase). The downstream steps were performed similarly to the analysis above. Pseudotime calculation and the assignment of milestones and segments were performed with setting two milestones (29: YS-derived macrophages, 2: monocytes) as roots (*scf.tl.roots*, *scf.tl.pseudotime*). To explore genes involved in the LYVE1+ macrophage maturation, the segments belonging to the trajectory (i.e. which formed a contiguous path) were subsampled, and the genes whose expressions were associated and upregulated along the pseudotime were selected (scf.tl.test association, sc.tl.rank genes groups). The pathways enrichment analysis was performed using the selected genes and GSEApy[60].

### 4.12 Cardiomyocyte gene signatures

The gene expression profile of SAN pacemaker cells (“SinoatrialNodeCardiomyocytes”) were compared to that of other cardiomyocytes and 675 DEGs were obtained (scanpy.tl.rank genes groups, logFC¿1, mean expression¿0.1). For the foetal-specific gene signatures, the foetal SAN pacemaker cell population was compared to the adult counterpart (SAN P cell[22]). For each cell type, donors were treated as replicates by making pseudo-bulk data (mean). Groups which have less than 10 cells were removed from the analysis. Wilcoxon rank-sum test was performed to identify DEGs (620 genes)(logFC¿1, mean expression¿0.1). Intersection of the two DEGs were used as foetal-specific pacemaker cell signatures (143 genes) in the downstream analysis. The gene signatures of the left ventricular cardiomyocyte populations were selected as follows. For compact or trabeculated cardiomyocyte-specific signatures, genes associated with and upregulated along the pseudofime (calculated per population) and expressed higher in one population compared to the other (either compact-vs-trabeculated or trabeculated-vs-compact) were selected (compact: 130 genes, trabeculated: 107 genes). For the common signature, genes associated with and upregulated along the pseudotime of both populations were selected (336 genes). Pathway enrichment analysis was performed for each gene signature using GSEApy[60].

### 4.13 Gene regulatory network analysis

The SCENIC+ pipeline [19] was used to predict TFs and putative target genes as well as regulatory genomic regions harbouring TF binding sites. For each analysis, gene expression and chromatin accessible region (peak) counts of multiome data was used as input. For the analysis of pacemaker cells, all the cardiomyocytes were used as input and metacells were created by clustering cells into groups of approximately 10 cells based on their RNA profiles and subsequent aggregation of counts and fragments. For the maturation analysis of atrial and left ventricle cardiomyocytes, the same multiome data used for the trajectory analysis were used as input. Metacells were created using SEACells[61](10 nuclei per metacell) based on the MultiVI latent space and subsequent aggregation of counts and fragments. For monocyte and macrophage populations, multiome data were subsetted from the data used for trajectory analysis, and metacells were not created because of the small nucleus number. CisTopic was applied to identify region topics and differentially accessible regions from the fragment counts as candidate regions for TF binding. CisTarget was then run to scan the regions for transcription factor binding sites and GRNBoost2 was used to link TFs and regions to target genes based on co-expression/accessibility. Enriched TF motifs in the regions linked to target genes were used to construct TF-region and TF-gene regulons. Finally, regulon activity scores (AUC) were computed with AUCell based on target-gene expression and target region accessibility. Networks of TFs, regions and target genes (eGRNs) were constructed by linking individual regulons. TF-region-gene links for all analyses can be found in Supplementary Table 3-5. Constructed network was further pruned for downstream analysis and visualisation. For the SAN pacemaker cells, GRNs which target SAN pacemaker cell gene signatures, ion channel genes, or axon guidance genes were first selected. For each TF-region-gene interaction, either TF (logFC¿1, mean expression¿0.1, proportion of expression¿0.1) or region (logFC¿0, proportion of accessibility¿0.01) being SAN pacemaker cell-specific (compared to other cardiomyocytes) were further selected. For the left ventricular cardiomyocytes, GRNs which target “compact” and “common” gene signatures were selected similarly to the GRNs of SAN pacemaker cells (compact cardiomyocyte GRNs). In-silico TF perturbation analysis was performed using CellOracle for the atrial cardiomyocyte and left ventricular cardiomyocyte analysis. CellOracle object was made using the genes included in the SCENIC+ regulons of each population. Input baseGRN was also prepared using the same regulon (only activator regulons). The cluster labels were generated by combining fine-grained cell types and binned chronological age (n bin = 3). TFs which govern the pruned GRNs were tested for in-silico perturbation.

### 4.14 Transmural axis analysis using OrganAxis

The spots which belong to left ventricle regions were manually selected (**Fig. S4G**). The inner (endocardial side) and outer (epicardial side) most layer were manually annotated. Using the OrganAxis algorithm[62], we calculated the relative distances of each spot from the two manually annotated layers. we calculated the L2 distance of all spots to the closest corresponding points in each annotation by number of KNN points (dist2cluster, kNN parameter = 4). To construct an axis anchored to the interface between two structures (“transmural axis”), we fitted a normalised signoidal curve to the relative distances of each spot and placed splots in relative positions in respect of the two structures (axis 2p norm). The gene expressions and cell type abundances were aggregated per bin along the transmural axis and clustered along the transmural axis and visualised (seaborn.clustermap). The cell type abundance was normalised for each individual sample (age) to account for potential technical variability related to different heart sizes. This normalisation enabled us to analyse the transitions of cell type localisations across the transmural axis. For gene selection, transmural axis was binned into three part (outer, middle, and inner) and differentially expressed genes for each part (compared to rest) were selected (logFC¿0.5, mean expression¿0.01). Pathway enrichment analysis was performed for the genes upregulated in outer or inner part. Nuclei segmentation of the H&E images of Visium sections was performed using Cellpose[63] implemented in Squidpy[64] (squidpy.im.segment, method=cellpose he, flow threshold=0.8, channel cellpose=1).

### 4.15 Cell-cell interaction analysis

For the analysis of sc/snRNA-seq data, we performed CellPhoneDB (v.5) analysis using all cells in the dataset to infer cell-cell interactions among fine-grained cell types: “statistical analysis method” with the default parameters and selected ligands and receptors, all of which were expressed in at least 5% of the cells. The ligand–receptor interactions were further selected on the basis of mean expression levels and the biological questions as indicated in the Results and the figure legends. For the analysis using Visium HD data (**Fig. 3J**), we performed CellPhoneDB (v.5) analysis utilising cell types and ligand-receptor genes, constrained by their spatial localisations within the identified SAN niche, as determined by TissueTypist. The “statistical analysis method” with the default parameters and selected ligands and receptors, all of which were expressed in at least 5% of the cells, were used. For plotting, the ligand-receptor interactions and the cell types were further selected on the biological questions as indicated in the Results and the figure legends.

### 4.16 smFISH

FFPE foetal heart tissue sections (5 *µ*m) were placed onto SuperFrost Plus slides. Staining was performed using the RNAscope Multiplex Fluorescent Reagent kit v2 assay (Advanced Cell Diagnostics, Bio-Techne), automated with a Leica BOND RX, according to the manufacturers’ instructions. Automated pretreatment included baking at 60 degree Celsius for 30 minutes, dewaxing, heat-induced epitope retrieval at 95 degree Celsius for 15 min in buffer ER2, and digestion with Protease III for 15 min. Following probe hybridisation, tyramide signal amplification with Opal 520 and Opal 650 (Akoya Biosciences) and TSA-biotin (TSA Plus Biotin Kit, Akoya) with streptavidin-conjugated Atto 425 (Sigma Aldrich) was used to develop RNAscope channels. All nuclei were DAPI stained. Stained sections were imaged using a Perkin Elmer Opera Phenix High-Content Screening System with a x20 water-immersion objective (NA of 0.16, 0.299 *µ*m per pixel). The following channels were used: DAPI (excitation, 375 nm; emission, 435-480 nm); Opal 520 (excitation, 488 nm; emission, 500-550 nm); Opal 650 (excitation, 640 nm; emission, 650-760 nm); and Atto 425 (excitation, 425 nm; emission, 463-501 nm).

### 4.17 Trisomy 21 single-nucleus multiome analysis

Post-quality control trisomy 21 (T21) single-nucleus RNA-seq data from four hearts (11–14 PCW) were further curated by removing non-cardiac cells and non-reproducible (across samples) cells. Coarse-grained and fine-grained cell type annotations were assigned using a label transfer approach (CellTypist), referencing euploid singlenucleus RNA-seq data. The T21 dataset was then integrated with age-matched euploid samples (12–14 PCW, three hearts), referred to as “Trisomy 21 + Euploid” in the associated web portal. Using the integrated dataset, the proportions of compact cardiomyocytes within the left and right ventricular cardiomyocyte populations were quantified. To generate a shared latent space for comparative analysis of T21 and euploid samples, and to assess T21-associated changes in cell population using the Milo[65], we performed donor-balanced subsampling across cell types (referred to as “Trisomy 21 + Euploid, subsampled for Milo analysis” in the web portal). Integration of the subsampled data was performed using scVI, accounting for donor identity and continuous covariates including total transcript counts, percentage of mitochondrial reads (%mito), and percentage of ribosomal reads (%ribo). The shared latent space generated by scVI was used to generate two dimensional UMAP embedding (**Fig. 5A**) and to construct a k-nearest neighbour graph (with n neighbors = 20), which formed the basis for identifying overlapping cellular neighbourhoods using milo.make nhood. Differential abundance testing was then conducted between T21 and euploid samples, and significantly altered neighbourhoods were identified (SpatialFDR ¡ 0.1). To perform differential gene expression analysis using cellular neighbourhoods as replicates, significantly enriched neighborhoods (FDR ¡ 0.1) associated with either T21 or euploid were used. For these neighbourhoods, we refined cell-to-neighbourhood assignments to ensure that each cell was uniquely assigned to a single neighbourhood, avoiding overlap across neighbourhoods. Gene expression data were then aggregated within each refined neighbourhood to generate pseudo-bulk expression profiles. These pseudo-bulk data were used as input for differential expression analysis using edgeR[66], comparing T21-enriched neighbourhoods to euploid-enriched neighbourhoods within each cell type. The resulting lists of differentially expressed genes were subjected to pathway enrichment analysis using gene set enrichment analysis (GSEA) using blitzgsea package[67] to identify pathways associated with T21-related transcriptional changes.

### 4.18 Mouse single-cell RNA sequencing

#### 4.18.1 Sample preparation and sequencing

Mice carrying the Dp(16Lipi-Zbtb21)1TybEmcf (Dp1Tyb) allele were described previously [35] and have been deposited with JAX (strain #037183). Dp1Tyb mice were backcrossed on a C57BL/6J genetic background for more than 10 generations. Mice were maintained in independently ventilated cages in a specific pathogen free facility and given food and water ad libitum. All animal experiments were carried out under the authority of a Project Licence granted by the UK Home Office and were approved by the Animal Welfare Ethical Review Body of the Francis Crick Institute. To generate embryos, Dp1Tyb female mice were crossed with C57BL/6J males. Hearts from 5 Dp1Tyb and 8 wild-type mouse embryos at day 13.5 of gestation (E13.5) were dissected and subsequently dissociated into single cells using the Neonatal Heart Dissociation kit and gentleMACS Octo dissociator (Miltenyi Biotec). Cells were resuspended in PBS, 0.04% BSA, filtered through 40 um FLowmi cell strainer and kept on ice until loaded. Approximately 20,000 cells were loaded on Chromium Chip and partitioned in nanolitre scale droplets using the Chromium Controller and Chromium. Next GEM Single Cell Reagents (CG000315 Chromium Single Cell 3’ Reagent Kits User Guide (v3.1 - Dual Index)). Within each droplet the cells were lysed and the RNA was reverse transcribed. All of the resulting cDNA within a droplet shared the same cell barcode. Illumina-compatible libraries were generated from the cDNA using Chromium Next GEM Single Cell library reagents in accordance with the manufacturer’s instructions (10x Genomics, CG000315 Chromium Single Cell 3’ Reagent Kits User Guide (v3.1 - Dual Index)). Final libraries are QC’d using the Agilent TapeStation and sequenced using the Illumina NovaSeq 6000. Sequencing read configuration: 28-10-10-90.

#### 4.18.2 Read mapping, data processing, and analysis

Each sample was mapped to the mouse reference genome (refdata-gex-mm10-2020-A) provided by 10x Genomics using the CellRanger software (CellRanger v 6.1.2) with default parameters. Downstream analysis was performed using the Scanpy package. Doublets with a score of ¿0.15 were removed using Scrublet. We performed quality control and filtering based on the following settings: number of genes¿200, % mitochondrial¡20 (cell), red blood cell score¡2. After pre-processing and removal of low-quality cells and nuclei, the coarse-grained and mid-grained cell types were annotated by transferring cell type labels from the human dataset (CellTypist). Single-cell transcriptomics data were integrated using scVI (Gayoso et al. 2022; Lopez et al. 2018) accounting for continuous covariates of total counts, %mito, and %ribo. Neighbourhood identification, high-resolution leiden clustering, and further dimensional reduction using UMAP were performed based on the scVI latent space. For differential abundance testing, overlapping cellular neighbourhoods were identified using Milo[65] and the calculated neighbourhood graph based on the scVI (n neighbors=10) latent spaces (milo.make nhoods). The differential abundance testing was performed between genotypes (wild-type and Dp1Tyb), and significantly enriched neighbourhoods were identified (SpatialFDR ¡ 0.25). Data has been deposited with the Gene Expression Omnibus (accession number: GSE240137). The differential gene expression analysis and GSEA analysis were performed similarly to the human T21 data, as described above.

### 4.19 Detection of apoptosis in iPSC-derived cardiomyocytes

#### 4.19.1 Stem cell lines and maintenance

This study used the trisomy 21 human induced pluripotent stem cell (hiPSC) line WC-24-02-DS-M and its isogenic euploid control WC-24-02-DS-B (WiCell Research institute). HiPSCs were maintained in feeder-free conditions on Geltrex^TM^ (Thermo Scientific) in mTESR1 (STEMCELL Technologies). Cells were passaged as aggregates every 3-4 days at a ratio 1:6 using a combination of enzymatic and mechanical dissociation. Karyotype and pluripotency were confirmed via the Karyostat+ and Pluritest assays (Thermo Fisher Scientific) respectively and genetic stability was monitored via routine low-pass sequencing (Genomics STP, Francis Crick Institute).

##### Cardiomyocyte differentiation

An adaptation of the standard WNT modulation protocol (PMID: 24930130, PMID: 23257984) was used to differentiate hiPSCs into cardiomyocytes. HiPSCs were dissociated to single cells and seeded on 12 well plates at a density of 5.7×10^4^ cells/cm^2^ in mTESR1 and 10µM Y-27632 (Tocris). Media was replaced daily with mTESR1 until cells reached 70-85% confluency (day 0), at which point media was switched to RPMI 1640 (Thermo Scientific) supplemented with B27-insulin (Thermo Scientific) and 12µM CHIR99021 (Selleck Chemicals). Media was replaced with RPMI+B27-insulin after 24 hours and then with RPMI+B27-insulin supplemented with 1µM IWR-1 (Sigma) and 65µg/ml L-ascorbic acid (Sigma Aldrich) after a further 24 hours. Routine media changes occurred every 48 hours thereafter with RPMI+B27-insluin + L-ascorbic acid until day 6, switching to RPMI+B27+L-ascorbic acid from day 8 onwards. Between days 10-13 cultures were subjected to a 72-hour period of ‘metabolic selection’ consisting of 24 hours’ incubation in DMEM(-glucose/-glutamine) (Thermo Scientific) followed by 48 hours of RPMI1640 without glucose + 4mM Lactate + L-ascorbic acid. Cardiomyocytes were re-plated routinely between days 14-16 and prior to relevant assays using a cardiomyocyte dissociation kit (STEMCELL Technologies) as per manufacturer’s guidelines.

##### TUNEL & Immunofluorescence Staining of hiPSC-derived cardiomyocytes

TUNEL staining was performed on hiPSC-derived cardiomyocytes at day 30-35 of differentiation. Prior to analysis iPSC-CMs were re-plated onto 96 well imaging plates (Greiner Bio-One) at a density of 4.5×10^4^cells/cm^2^ and allowed to adhere. Cells were fixed with 4% PFA for 15 minutes at room temperature. TUNEL staining to detect apoptotic cells was performed using the Click-iT™ Plus TUNEL Assay (Invitrogen) as per manufacturer’s instructions. Subsequently samples were stained with mouse anti-cardiac troponin T (Invitrogen, MA5-12960) overnight at 4°C, followed by donkey-anti-mouse-568 secondary antibody (Invitrogen, A10037) for 60 minutes at room temperature and DAPI (Sigma-Aldrich) for 5 minutes at room temperature. Cells were imaged using the Operetta CLS high-content analysis system (PerkinElmer) and analysis of TUNEL positive cardiomyocytes (cardiac troponin T positive cells) was performed using the Harmony 5.2 software. Nuclei were segmented based on DAPI staining and cardiomyocytes were identified using a threshold for cardiac troponin T (Alexa568) intensity in the cell area surrounding but not including the nucleus (‘surrounding region’). TUNEL positive cells were identified using a threshold for TUNEL stain (Alexa 647) intensity within the nucleus region. Cells meeting the threshold for both ‘cardiomyocyte’ and ‘TUNEL positive nuclei’ were identified and quantified as TUNEL positive cardiomyocytes.

### 4.20 Data availability

Open-access datasets will be available from ArrayExpress (www.ebi.ac.uk/arrayexpress) with accession numbers (Multiome snATAC-seq and Visium). All of the processed data are available for browsing gene expression and download from the Heart Cell Atlas (https://www.heartcellatlas.org/foetal.html, currently password protected and will be publicly available at the time of publication). Inferred gene regulatory networks are available in Supplementary Table 3-5. The human reference genome (GRCh38) used for read mapping is available from 10x Genomics (https://support.10xgenomics.com/single-cell-gene-expression/software/release-notes/build). The mouse scRNA-seq data has been deposited with the Gene Expression Omnibus (accession number: GSE240137, will be publicly available at the time of publication).

### 4.21 Code availability

The code used for all the analysis performed in this study is available on GitHub at https://github.com/Teichlab/Developing_heart_atlas. The TissueTypist Python package is available at GitHub (https://github.com/Teichlab/TissueTypist).

## 5 Acknowledgements

We thank the donors for granting access to the tissue samples. We also thank staff at the Wellcome Sanger Cytometry Core Facility, Cellular Genetics Informatics team, Spatial Genomics Platform team, and Core DNA Pipelines team for their support; A.Oszlanczi for her help on sample management; B. Cakir and J. Cox for their help on the Heart Cell Atlas web portal; and A.Wilk for administrative assistance; R. Barker for tissue procurement.

Illustrations in Figs. 1A and 3K were created using BioRender (https://biorender.com). This work was made possible by a partnership between the Wellcome Sanger Institute and Cambridge Stem Cell Institute at University of Cambridge.

This project was made possible in part by the Wellcome Trust (WT206194 to S.A.T.); the Chan Zuckerberg Foundation (2021-237882 to S.A.T.); BBSRC strategic Longer Larger grant (to S.A.T., S.S. and R.T.); S.A.T. is also funded by the CIFAR Macmillan Multi-scale Human Program; V.L.J.T. was supported by The Francis Crick Institute which receives its core funding from Cancer Research UK (CC2080), the UK Medical Research Council (CC2080), and the Wellcome Trust (CC2080); the Wellcome Trust Clinical PhD Fellowship to J.C.; the Overseas Research Fellowship of the Takeda Science Foundation to K.K.; the British Heart Foundation (BHF) Senior Fellowship (FS/18/46/33663) (S.S. and L.G.); the Oxbridge BHF Centre for Regenerative Medicine (RM/17/2/33380)(V.K.S.); and BHF grants PG/17/24/32886 (L.G.) and RG/17/5/32936 (H.D.); This project has received funding from the European Union’s Horizon 2020 research and innovation programme under the Marie-Sk-lodowska-Curie grant agreement No. 101026233 to J.P.P.; We also acknowledge core support from the Wellcome Trust, the Medical Research Council and the Wellcome Trust–Medical Research Council Cambridge Stem Cell Institute. This research was funded, in whole or in part, by the Wellcome Trust (grant no. 203151/Z/16/Z).

## 6 Author contributions

Project design, Data generation, Analysis, Manuscript: J.C., K.K. Project design, Analysis, Manuscript: S.B., V.K.S. Data generation (Trisomy 21 iPSC-cardiomyocytes): R.H. Data generation (Dp1Tyb scRNA-seq, Trisomy 21 tissue procurement): E.L.E., R.A. Analysis (GRN): J.P.P., J.P. Data generation (Immunofluorescence): A.W.C. Analysis (Visium mapping): K.P. Data generation (Visium): M.D., I.M., H.J., M.P. Data generation (Multiome): L.R., C.I.S., R.K., S.P. Data generation (tissue procurement): X.H. Analysis (Tissue structure annotation): S.Y.H. Analysis (OrganAxis): N.Y. Data generation (smFISH): L.T., K.R. Heart collection, processing and sequencing: H.D., L.G. Guide on the experiment and analysis, reviewing/editing manuscript: R.T., A.B., V.T. Supervision: K.K., S.S., S.A.T.

## 8 Supplementary Figures

**Supplementary Figure S1.**
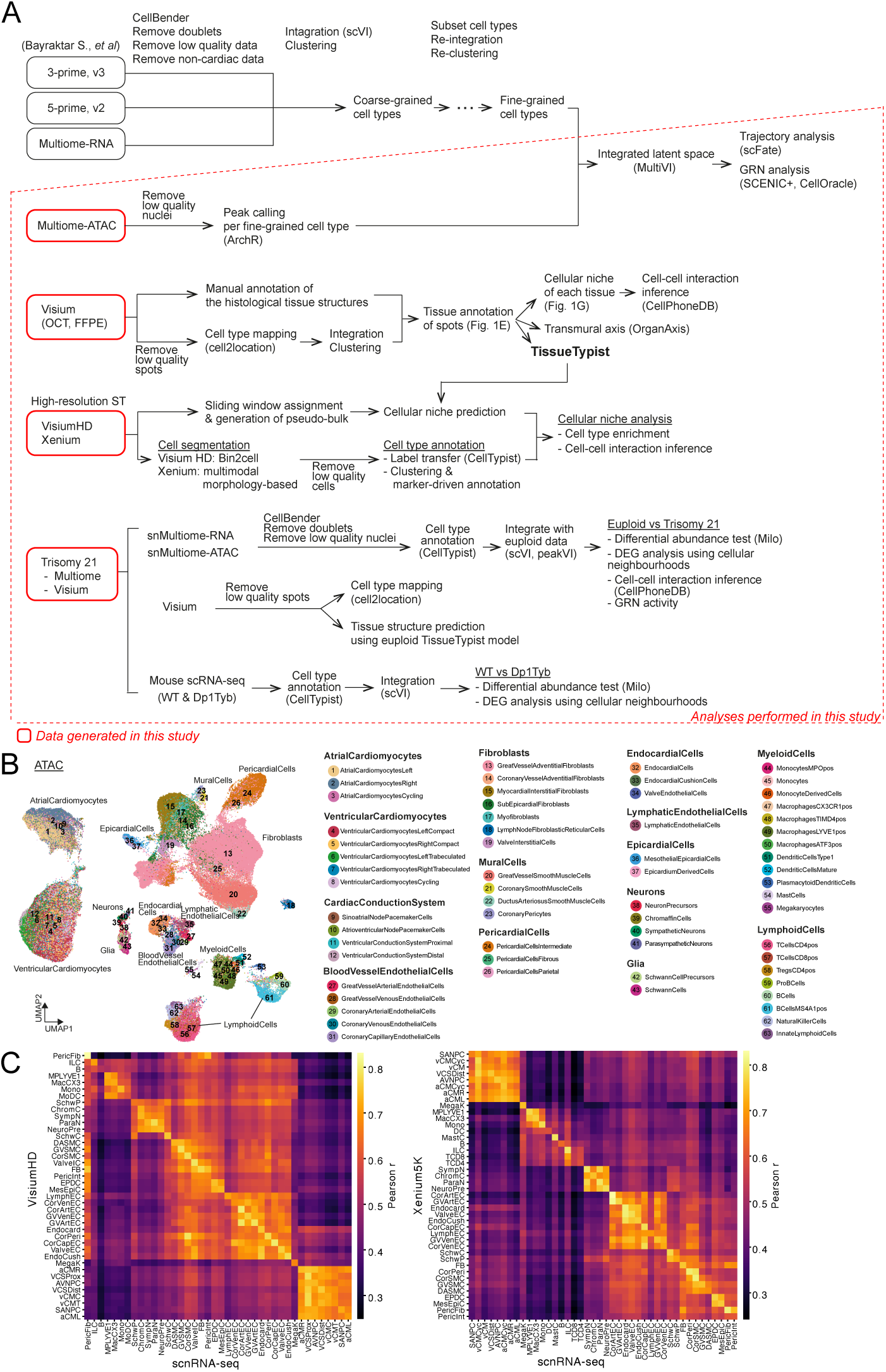
**A:** Workflow of the data processing and analyses performed in this study. **B:** UMAP embedding of the chromatin accessibility data labelled with the fine-grained cell types. 63 fine-grained cell type annotations are provided adjacent to the embeddings, grouped under their respective mid-grained cell types. **C:** Pearson’s collerations of gene expressions per fine-grained cell type between the Visium HD (left) or Xenium5K (right) data and scnRNA-seq datasets. *Abbreviations*: The abbreviations of the fine-grained cell types are listed in Supplementary Table 6.

**Fig. S2:**
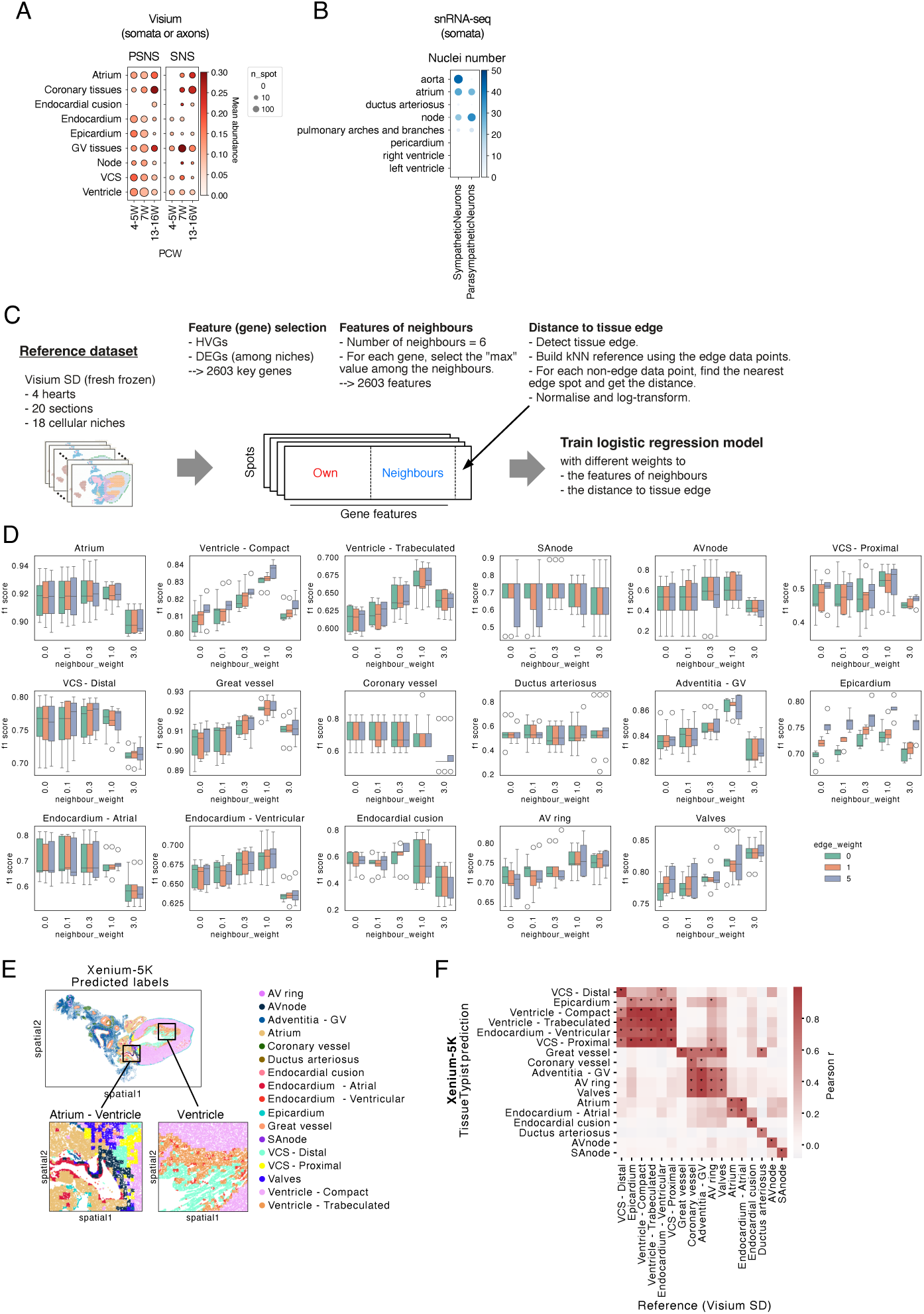
**A:** Dot plot showing estimated cell abundance (cell2location) of parasympathetic neurons (PSNS) or sympathetic neurons (SNS) along age in the Visium data. The colour scale represents the mean cell abundance, and the dot size represents the number of spots which include PSNS or SNS (abundance threshold=0.05) in each annotated tissue and age bin as indicated. This result includes localisat5i9ons of both somata and axons of the neuronal populations. **B:** Dot plot showing nuclei number of sympathetic or parasympathetic neurons in each anatomical region. The snRNA-seq data with the anatomical regions dissected was used for this analysis (14, 15, and 20 PCW hearts). The colour scale and the dot size represent the nuclei number. This result indicates localisations of somata. **C:** The workflow of the TissueTypist model training. **D:** Cross-validated F1 scores for the tissue structures across different weightings of neighbouring-spot information and distance-to-edge features. **E**: Prediction results on Xenium-5K data of 9 PCW heart. **F**: Correlations of cell-type proportions per niche between the query Xenium-5K data and reference datasets.

**Supplementary Figure S3.**
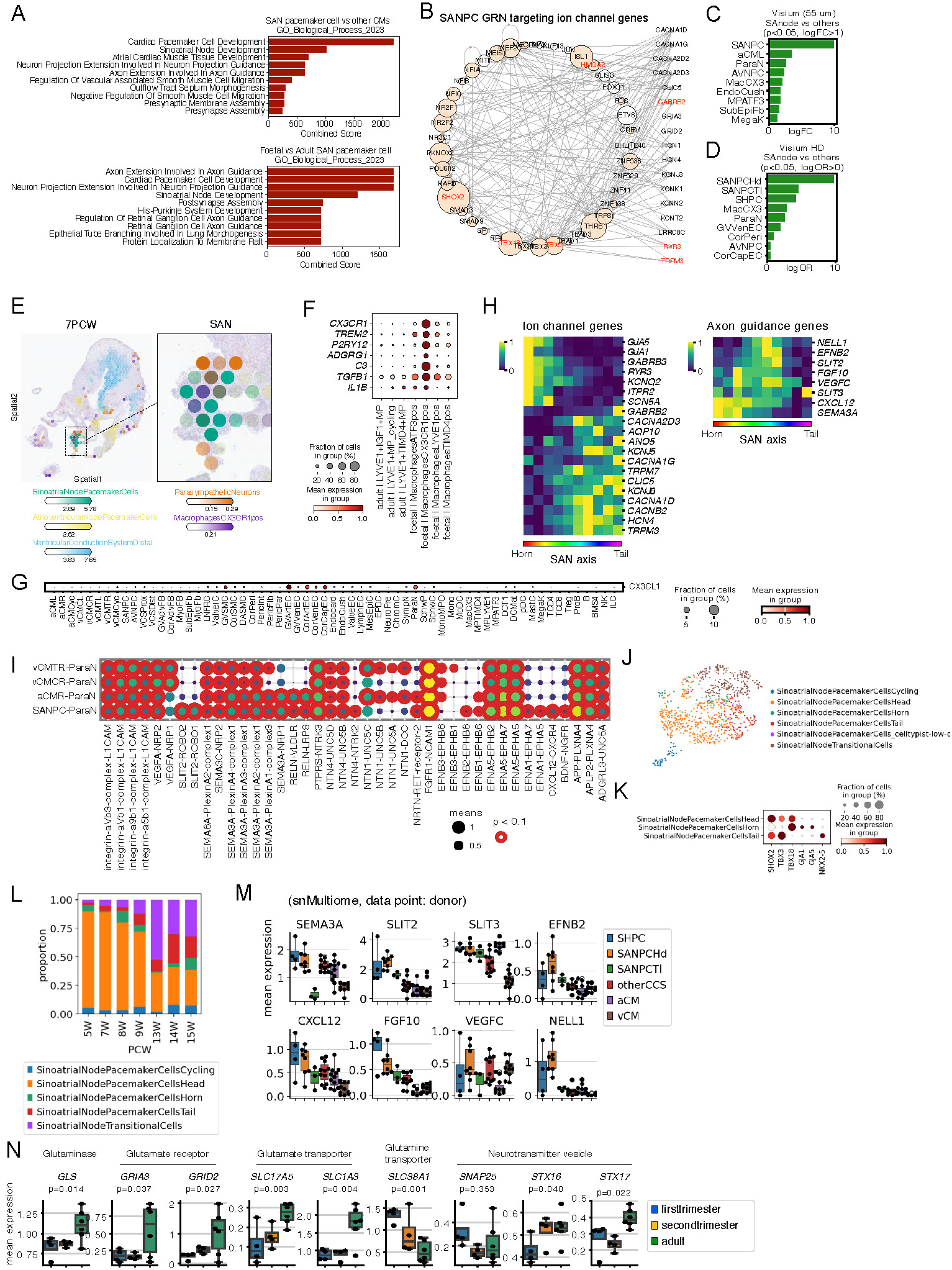
**A:** Pathway enrichment analysis (GO, biological process) of the genes highly expressed in the foetal SAN pacemaker cells compared to the other foetal atrial cardiomyocytes (top) or the genes narrowed down by comparing to the adult SAN pacemaker cells (bottom). **B:** Gene regulatory network (GRN) targeting ion channel genes specifically expressed in SAN pacemaker cells. Transcription factors (TFs) are shown in a circular arrangement with ion channel target genes (TGs) on the right. The edges represent TF-TG activation. The size of the TF nodes represents the centrality score (eigenvector) in the TF network. TFs, which directly interact with the target genes, are coloured by salmon. Genes in red are expressed higher in foetal pacemaker cells compared to their adult counterpart. **C, D:** Bar plots showing cell types significantly enriched in the SAN compared to the other tissues. Using the Visium (donor C83, 3 sections)(C) or Visium HD (donor C194, 1 section)(D) data. **E:** Estimated cell abundance of the indicated cell types is projected on the 7 PCW heart (Visium). Colocation of pacemaker cells, parasympathetic neurons, and CX3CR1+ macrophages at the SAN is highlighted. **F:** Dot plot showing expression of microglial genes involved in neuronal development and homeostasis, comparing adult and foetal cardiac macrophage populations. **G:** Dot plot showing the *CX3CL1* expression across the fine-grained cell types from the all time points. **H:** Heatmap showing scaled gene expression (min-max normalisation per gene) of ion channel genes (left) or axon-guidance genes (right) along the SAN axis. **I:** Inferred cell-cell interactions involved in axon guidance between the cardiomyocytes and the parasympathetic neurons (ParaN) in the SAN niche. The colour scale and dot size represent the mean expression levels of the interacting ligand-receptor partners. Cell type-specific interactions are delineated with a red border. **J:** UMAP embedding of the single-nucleus Multiome data (gene expression) of the SAN pacemaker cells. The subpopulations were annotated using a trained label-prediction model (CellTypist) trained on the Visium HD pacemaker cell annotations (Fig. 3F). **K:** Dot plot of the single-nucleus Multiome data showing the expression of maker genes for pacemaker cell populations in the sinus horn, SAN-Head and SAN-Tail. **L:** Bar plot illustrating the proportion of pacemaker subpopulations across different ages. **M:** Expression of the genes involved in axon guidance in the SAN pacemaker cell populations and other CMs. The dots show the mean expression of each donor. For the box plots, the centre line shows the median, the box limits represent the 25th and 75th percentiles, and the whiskers show the minimum and maximum values. **N:** Expression of the genes involved in glutamate signalling in the SAN pacemaker cells are shown. The dots show the mean expression of each donor. For the box plots, the centre line shows the median, the box limits represent the 25th and 75th percentiles, and the whiskers show the minimum and maximum values. Statistical significance was assessed using a linear regression model across ordered stages (first trimester, second trimester, and adult), with p-values provided. **Abbreviations**: SAN – sinoatrial node; SANPC – SAN pacemaker cell; CM – cardiomyocyte; SHPC – sinus-horn pacemaker cell; ParaN – parasympathetic neuron; aCMR – right-atrial cardiomyocyte; Additional fine-grained cell-type abbreviations appear in Supplementary Table 6.

**Supplementary Figure S4.**
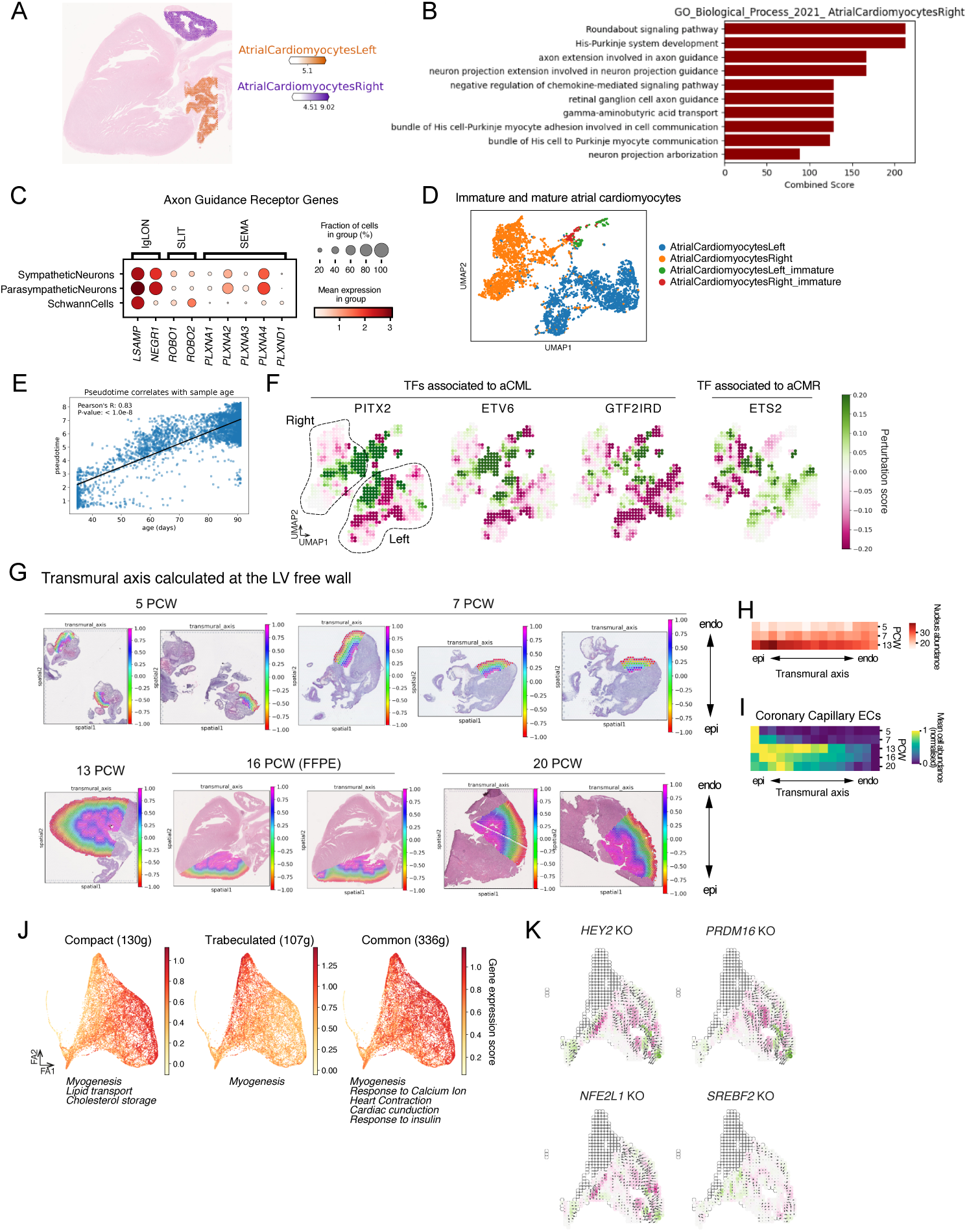
**A:** Cell2location mapping of L and R atrial cardiomyocyte signatures in a 16PCW heart. **B:** Bar plot showing pathway enrichment analysis result for the genes upregulated in RaCMs compared to LaCMs. Threshold set at log fold change ¿ 2, adjusted p-values ¡0.05, method: t-test. **C:** Dot plot showing mean expression of receptors for the ligands in Fig. 4D in neural cell types. **D:** UMAP embedding (multiVI space) showing annotation of immature atrial cardiomyocytes (these signatures were used for cell2location analysis in Fig. 4C. **E:** Scatterplot showing correlation between chronological age in day (smoothed across neighbourhood graph) and pseudotime. **F:** Representative plots of TF perturbation simulation results. Showing perturbation scores for impact on the maturation of atrial cardiomyocytes overlaid on the UMAP embedding. **G:** Assignment of transmural axis for the Visium spots on the left ventricle region. **H, I:** Normalised cell abundance (I) or nuclei number per spot (H) along the left ventricular transmural axis and the chronological age. The transmural axis was binned and the mean of the values were calculated for each bin and age. **J:** Compact-enriched (130 genes), trabeculated-enriched (107 genes) and commonly shared (336 genes) signatures associated with the pseudotime projected on the force-directed graph (Fig. 4K, L). Pathways identified by gene set enrichment analysis are italicised. **K:** Representative plots of TF perturbation simulation results. Showing perturbation scores for impact on the maturation of compact cardiomyocytes overlaid on the compact cardiomyocyte regions of the UMAP embedding (trabeculated cardiomyocyte regions are intentionally blank).

**Supplementary Figure S5.**
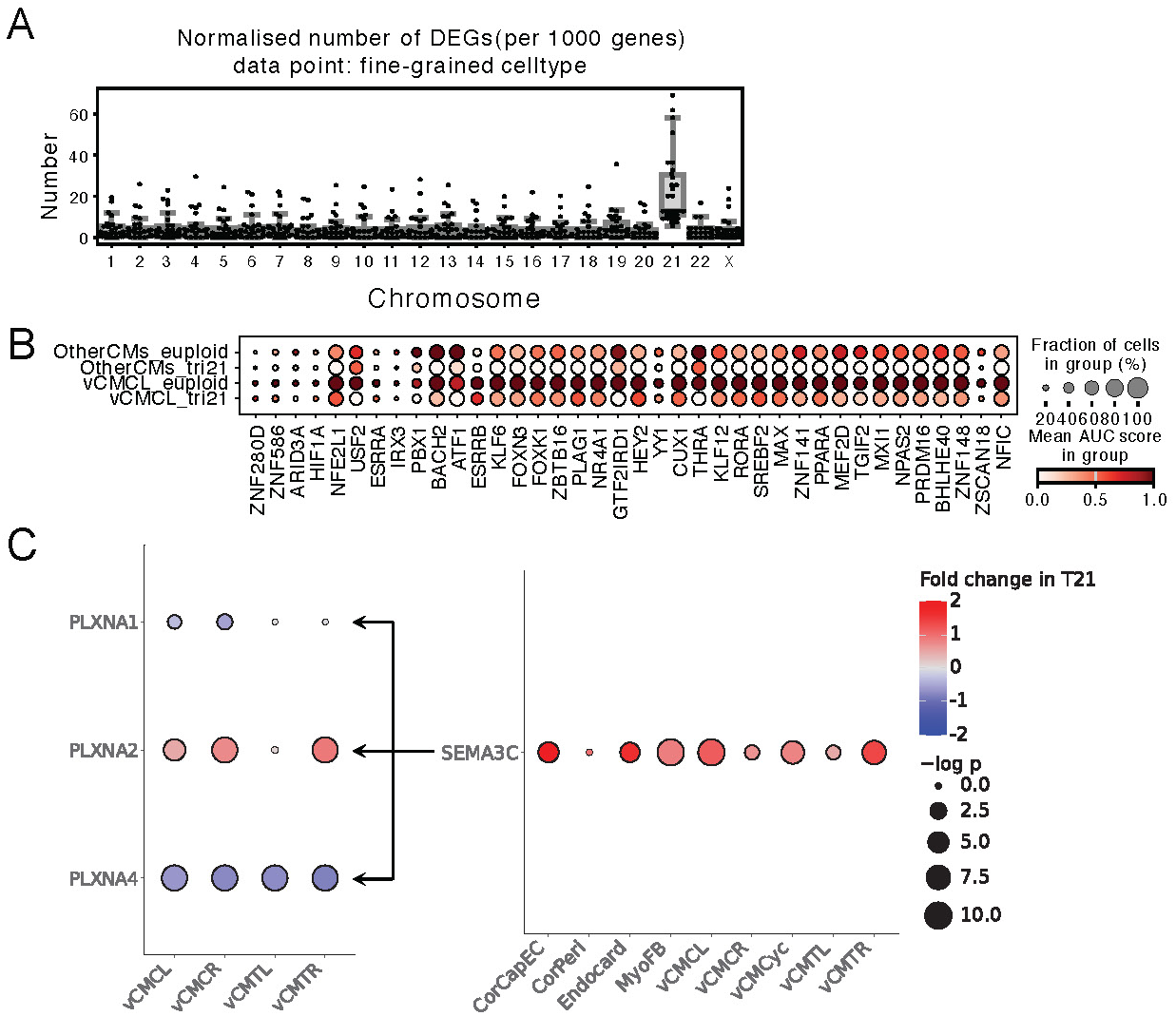
**A:** Box plot illustrating the normalised number of differentially expressed genes (DEGs) between Trisomy 21 and euploid samples, categorized by fine-grained cell type for each chromosome. The DEGs are normalised to a count of 1000 for comparison across different lengths of chromosomes. **B:** Dot plot showing the regulon activity scores (AUC score) of compact cardiomyocyte GRNs, whose scores were reduced in the T21 left compact cardiomyocytes compared to the euploid counterpart (logFC ¡ -0.1). **C:** Inferred cell-cell interactions mediated by Semaphorin-Plexin signalling between ventricular cardiomyocytes (vCMCL, vCMCR, vCMTL, vCMTR) and other cell types in the ventricular niche. The color scale indicates the fold-change in gene expression observed in Trisomy 21, while the dot size represents the statistical significance of the differential expressions. Abbreviations: The abbreviations of the fine-grained cell types are listed in Supplementary Table 6.

**Supplementary Figure S6.**
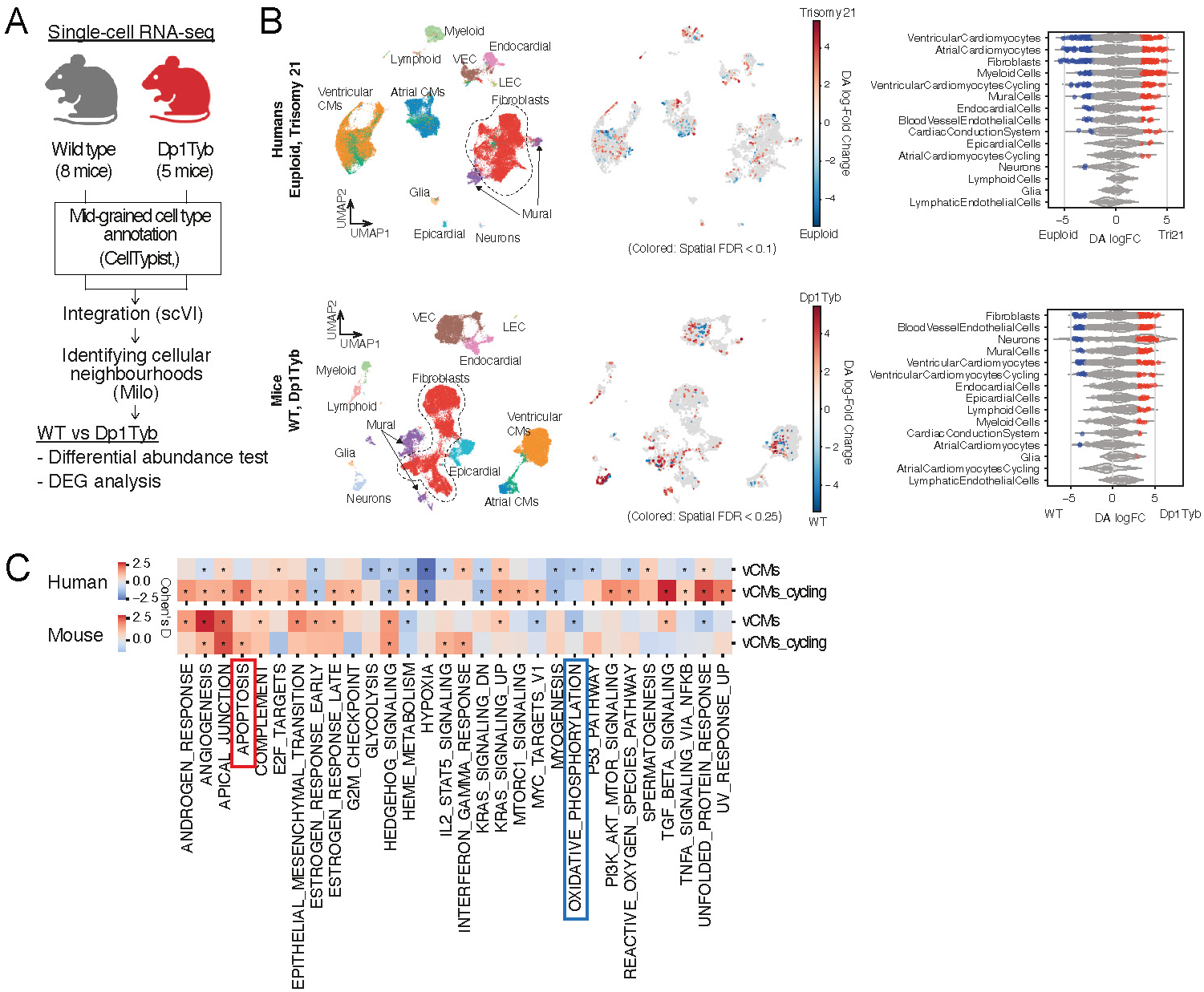
**A:** Schematic illustrating the mouse single-cell RNA-seq data analysis. **B:** UMAP embedding and differential abundance analysis (Milo) for human singlenucleus Multiome (gene expression) (top) and mouse single-cell RNA-seq (bottom) data. The bee-swarm plot depicts log-fold change (x-axis) in cell abundance between euploid and T21 or WT and Dp1Tyb mouse (scVI latent space), categorised by mid-grained cell type. Neighbourhoods overlapping the same cell type are grouped (y-axis). Neighbourhoods with significant differential abundance are coloured red or blue (Spatial FDR¡0.1 or 0.25). **C:** Heatmap illustrating the results of Gene Set Enrichment Analysis (GSEA) derived from differential gene expression analysis utilising cellular neighbourhoods (Milo). The assignment of neighbourhoods has been refined to eliminate the duplication of cells across multiple neighbourhoods. The colour scale indicates the normalised enrichment score associated with Trisomy 21.

### 8.1 Epigenetic stability of cells in the developing heart

The impact of cis-regulatory elements and epigenetic regulation on gene expression during cardiac development and cell type specification is significant [**? ?**]. Integrating single-nucleus RNA- and ATAC-seq data enables us to explore epigenetic stability, indicative of the chromatin status dynamics. To investigate this, we initially defined overlapping cellular neighbourhoods [36] in both the RNA and ATAC spaces. For each cellular neighbourhood identified in the RNA space, the degree of dispersion of the corresponding nuclei distribution across the cellular neighbourhoods identified in the ATAC space was calculated (entropy score) (**Fig. S7A**). In the cardiomyocyte populations, working cardiomyocytes exhibited elevated entropy scores (indicating reduced epigenetic stability) relative to cardiac conduction system cells, such as sinoatrial pacemaker cells or distal ventricular conduction system cells (**Fig. S7B, C**). This suggests that CCS cells have more stable and established epigenetic status, whereas working cardiomyocytes are undergoing chromatin remodelling. The entropy scores of monocyte and macrophage populations were highest among the myeloid cell types, potentially indicating their enhanced reactivity and dynamic development, influenced by the tissue environment and internal stimuli (**Fig. S7B, C**). In heart failure, monocytes and macrophages exhibit remarkable epigenetic plasticity, which facilitates their adaptive responses. This plasticity contributes to greater diversity and enhanced polarisation among macrophages[68]. Generally, embryonic macrophages are regarded as highly adaptable immune cells with considerable plasticity, which enhances their potential to regulate various biological processes[69]. It is likely that this characteristic enables them to seamlessly integrate into the cardiac microenvironment. It is important to note that these results may reflect the temporal dynamics of differentiation – a cell type that is still undergoing differentiation might be expected to display a wider collection of chromatin states relative to another cell type which has completed its differentiation process.

**Supplementary Figure S7.**
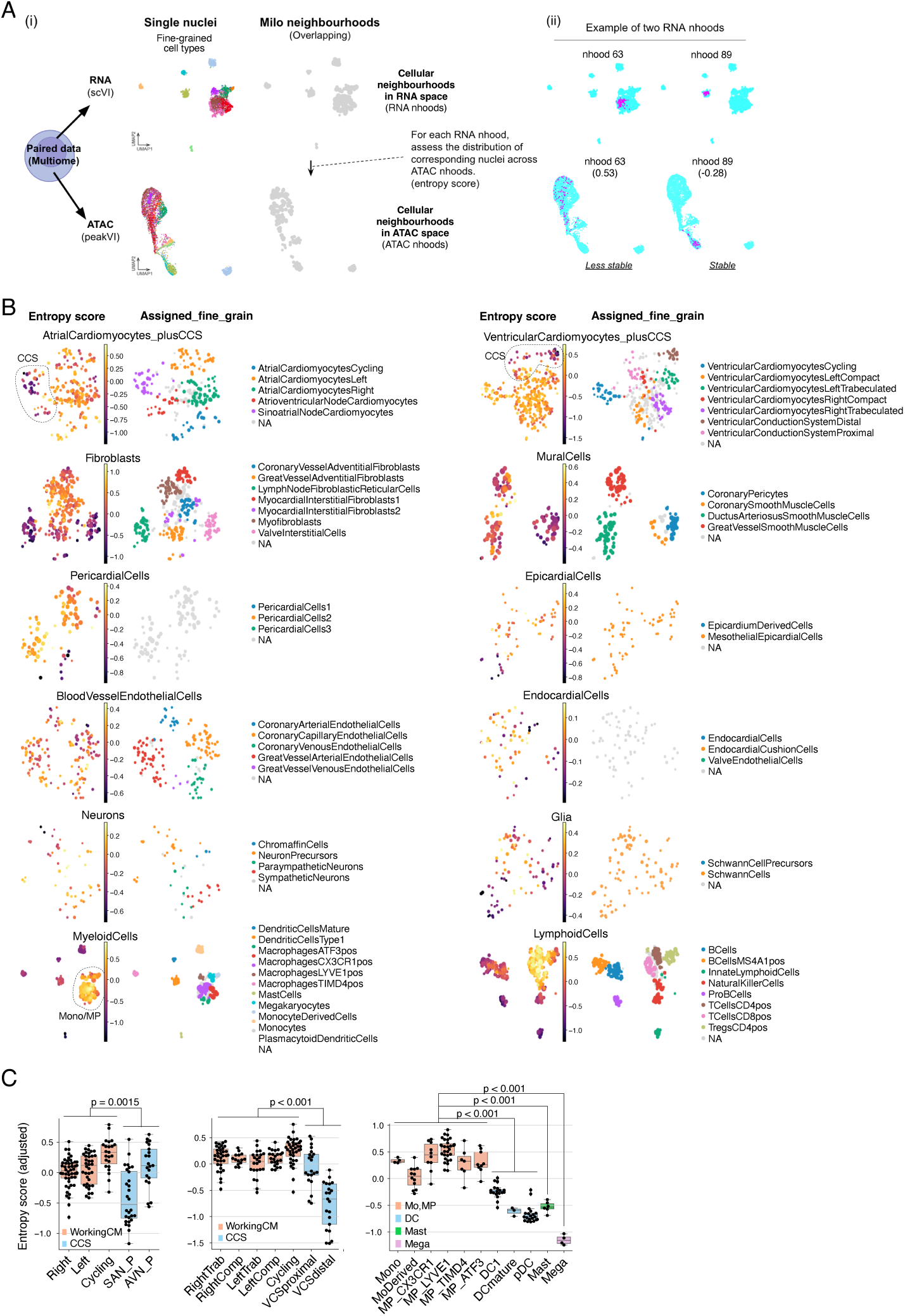
**A:** (i) The workflow of the epigenetic stability assessment. Using Multiome data, for each mid-grained cell type, the overlapping neighbourhoods (Milo) were found using gene expression data and scVI latent space (RNA neighbourhoods) or chromatin accessibility data and peakVI latet space (ATAC neighbourhoods). For each RNA neighbourhood, the entropy of the distribution of corresponding nuclei across the ATAC neighbourhoods was calculated. Data points represent neighbourhoods. (ii) Example distribution of two RNA neighbourhoods in the ATAC space (UMAP embedding based on peakVI latent space) are shown. Data points represent cells. RNA neighbourhood-63 shows greater scattered distribution in the ATAC space and a higher entropy score (0.53), suggesting that the neighbourhood cells are epigenetically unstable and potentially in the process of transitioning to another state. **B:** Entropy score (plot on the left of each mid-grained cell type) or assigned fine-grained cell type labels (plot on the right) projected on the RNA space (UMAP embedding based on scVI latent space). Data points represent RNA neighbourhoods, and the dot size shows the neighbourhood size. Cellular neighborhoods with a purity exceeding 0.6 are assigned the most abundant cell type. Neighbourhoods with no cell type exceeding this threshold are marked as NA. **C:** Box plots indicate the entropy scores of the neighbourhoods in each assigned cell type label of atrial or ventricular cardiomyocytes or myeloid cells. Data points represent neighbourhoods. *p*-values are provided for the comparison among groups of cell types as indicated (two-sided Kruskal-Wallis test, for multiple comparisons, post hoc analysis with Dunn’s test was performed with adjusting p-values using the Bonferroni method).

### 8.2 Cardiac macrophage development

Macrophages are critical in the embryonic development of various tissues^20^ including the heart^21^. We utilised gene expression and chromatin accessibility data of macrophages and monocytes and created an integrated embedding using multiVI^22^ (**Fig. S8A**). The principal graph was calculated based on the multiVI representation, and the pseudotime was obtained by setting the youngest milestone as the root (**Fig. S8B,C**). Recently, we described monocyte-independent macrophage differentiation in the human yolk sac (YS)^23^ that seeds developing organs. To test whether the embryonic heart contains macrophages originating from this route, we trained a CellTypist model with the YS dataset and transferred the labels to our dataset. The population found in the youngest milestone (5̃PCW) was predicted to be YS premacrophages (YSpMPs) with high probability (MacrophagesYSderivedLike)(**Fig. S8D**). High activities of the YSpMP regulons (FLI1, MEF2C)[70] at this population also suggest that the YSpMPs populate the early-stage heart (**Fig. S8E**). We also found that CX3CR1^+^ macrophages were predicted as a “microglia-like” population using the YS CellTypist model and expressed microglia marker genes (**Fig. S8D**), including *CX3CR1*, *TREM2*, *P2RY12*, *ADGRG1* and *C3* (**Fig. S8F**). We showed this microglia-like macrophages localise in the SAN (**Fig. 2A**, **Fig. 3D**, **Fig. S3C-E**) and express genes encoding ligands and receptors involved in synaptic maintenance (**Fig. S8G**)[71]. Recently it was shown that CX3CR1+ macrophages populate tissues outside the CNS, including skin, testicles, heart and aorta, during human development [72]. However, their precise localisation and function in the developing heart were not clarified. This finding suggests they are part of the developing sinoatrial node niche. Using the adult heart cell atlas, macrophage populations do not exhibit expression of *CX3CR1* or any other markers typically associated with microglia (**Fig. S3G**), suggesting their specific role in the developing heart.

For the LYVE1^+^ macrophage population, the chronological age of the trajectory from the MacrophagesYSderivedLike (**Fig. S8C**, red arrow) was correlated with the calculated pseudotime (**Fig. S8H**), reinforcing the notion that the pseudotime captures the maturation process. The top MSigDB Hallmark^24^ pathway enriched in the pseudotime-associated receptor ligand genes was angiogenesis, containing genes such as *VEGFA*, *SPP1*, *APP* and *CXCL8* (**Fig. S8I**). Spatial transcriptomic analysis showed colocalisation of LYVE1+ macrophages and vessels, such as coronary and lymphatic vessels (**Fig. 2A**, **Fig. S8J**), consistent with a previous study in mice^25^. We validated their localisation in coronary vessels using multiplex smFISH (RNAscope) (**Fig. S8K**). Finally, cell-cell interaction analysis revealed potential ligand-receptor interactions, suggesting that these LYVE1+ macrophages may facilitate angiogenesis through paracrine communication with smooth muscle and endothelial cells, including VEGFA signalling^26^ (**Fig. S8L**). These results bring new clarity to the emergence and role of cardiac macrophages in the developing heart.

**Supplementary Figure S8.**
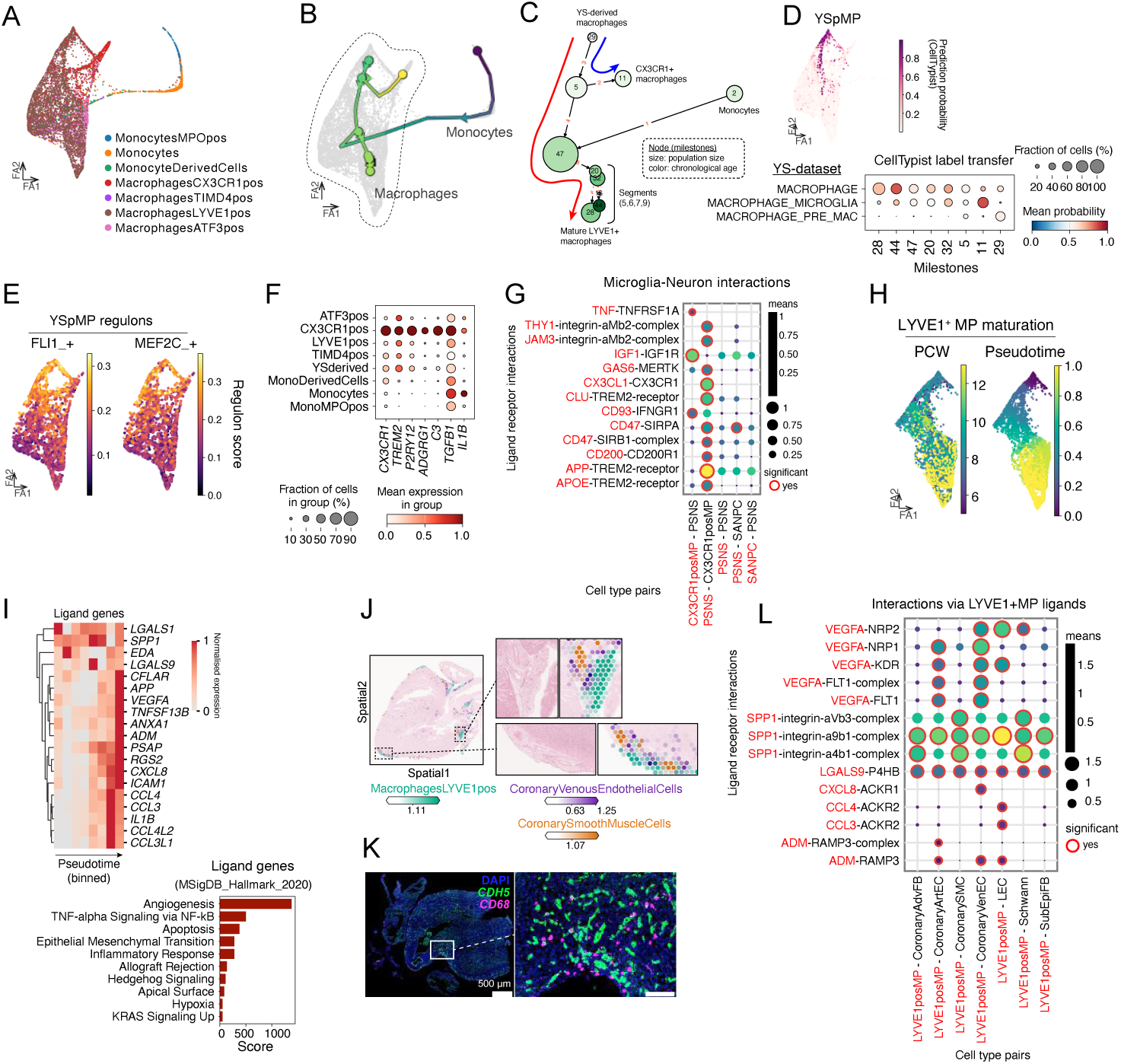
**A, B:** Force-directed graph of macrophages and monocytes using MultiVI latent space (based on both gene expression and chromatin accessibility). Monocyte and macrophage fine-grained cell types (A) or the principal graph coloured with pseudotime (B) are projected on the embedding. **C:** Graph showing macrophage and monocyte trajectory milestones and segments defined using MultiVI latent space and scFate workflow. The trajectories of LYVE1+ (red) and CX3CR1+ (blue) macrophage maturation are highlighted with arrows. The youngest milestone of macrophage (milestone 29) and monocyte (milestone 2) were set as the roots for the pseudotime calculation. **D:** Label transferring of yolk-sac (YS) macrophages to the foetal heart monocytes and macrophages was performed using CellTypist. The prediction probability of YS premacrophage (YSpMP) is projected on the UMAP. Dot plot showing mean probability and proportion of CellTypist prediction results for each macrophage. YS macrophage dataset [70] was used for the model training. **E:** Calculated activity scores of the YS premacrophages regulons (FLI1 and MEF2C)[70] are projected on the Force-directed graph. **F:** Dot plot showing the expression of maker genes for CX3CR1-positive microglia-like macrophages. **G:** Inferred cell-cell interactions of the SAN niche. The interactions are selected based on the ligands and receptors known for neuron-microglia crosstalk. Colour scale and dot size represent the mean expression levels of the interacting ligand-receptor partners. Cell-type specific interactions are delineated with a red border. **H:** Cells and nuclei belong to the segments associated with the LYVE1+ macrophage maturation (**Fig. S7C**, red arrow). The colour scales show chronological age (PCW, left embedding) or normalised pseudotime (right embedding). **I:** Heatmap shows the normalised gene expression of the pseudotime-associated ligand genes (HGNC Group ID: 542). The pseudotime was binned and the mean of the values were calculated for each bin. Bar plot shows the result of pathway enrichment analysis of the genes associated with the LYVE1+ macrophage maturation trajectory. **J:** Estimated cell abundance of LYVE1+ macrophages and coronary vessel cells (cell2location). The colocation of the cell types is highlighted. **K:** Multiplex smFISH (RNAscope) of the 7 PCW foetal heart region for *CD68* (yellow) and *CDH5* (red). DAPI was used to stain nuclei. **L:** Inferred cell-cell interactions between LYVE1+ macrophages and the other cell types in the coronary vessel niches. Colour scale and dot size represent the mean expression levels of the interacting ligand-receptor partners. Cell-type specific interactions are delineated with a red border.

